# Projective LDDMM: Spatially Reconstructing a Story of Rostrally-Dominant Tau in Alzheimer’s Disease

**DOI:** 10.1101/2022.03.16.484623

**Authors:** Kaitlin Stouffer, Menno Witter, Claire Chen, Eileen Xu, Marilyn Albert, Susumu Mori, Juan Troncoso, Daniel Tward, Michael Miller

**Author notes:** Contributing authors. These authors contributed equally to this work (Daniel Tward, Michael Miller).

## Abstract

Since Braak’s initial histological observations, it has been recognized that Alzheimer’s disease (AD) neurofibrillary tangles (NFTs) appear in the medial temporal lobe (MTL) of the brain very early in the disease course. MRI-based shape diffeomorphometry markers have demonstrated pre-clinical AD changes in the MTL but it has not been possible to confirm that these MRI changes correspond to the presence of NFTs. Here, we present a method termed Projective LDDMM for aligning sparse measurement profiles of AD pathology (i.e., 2D digital histology images) with 3D MRI. We reconstruct measures of 2D NFT density in the dense metric of 3D MRI, using the Mai Paxinos Atlas coordinates for two cases of advanced AD. Analyses reveal the highest levels of NFT density in the rostral third (10-15 mm) of the hippocampus and the adjoining regions of the entorhinal cortex and amygdala. These findings emphasize the selective vulnerability of MTL subregions in AD, and suggest that high resolution MRI methods might benefit from focusing on the rostral MTL to more closely link these MRI images to AD neuropathology.

---

Alzheimer’s disease (AD) is the leading cause of dementia worldwide [1]. Diagnosis and characterization of AD in its *early* stages remain key challenges, as existing technologies limit the identification of the neuropathological patterns thought to emerge years before symptom onset [2, 3, 4]. In clinical practice, AD is typically first characterized by progressive clinical changes in memory and behavior, and subsequently through imaging changes that indirectly reflect AD neuropathology (i.e. misfolded proteins, tau and amyloid-Beta (A*β*)) [5, 6, 7]. Efforts to identify and understand the spatiotemporal profile of AD in its early stages have centered on these biomarkers [8]–measures that indirectly reflect the underlying pathology, which are obtainable over the course of disease. Of the methods used, neuroimaging has emerged as a prominent player with the ability to localize pathology non-invasively (e.g. tau/amyloid PET) [9, 10], and with proposed surrogates such as shape diffeomorphometric markers (e.g. MRI) [11, 12, 13]. While these imaging measures have shown consistency with Braak staging [5, 6], an accurate rendering of the 3D spatiotemporal profile of tau and A*β* at the micron scale has not been achieved [9, 10]. The principle challenge has been integrating the 2D sparse measurements of histology, which are direct measures of disease, to the MRI 3D markers which are at much lower in-plane resolution.

This paper focuses on a new class of image-based diffeomorphometry methods which we term Projective LDDMM for aligning sparse 2D histological profiles to 3D coordinate systems across micron and millimeter scales. The 3D MRI to 2D digital histology mapping is representative of a class of multiscale, multi-modality mapping in biomedical research including traditional light microscopy mapping to dense reference atlases [14, 15, 16, 17], light sheet methods [18, 19], and spatial transcriptomics [20, 21, 22, 23]. We formulate the dense mapping of atlases to sparse images problem using the random orbit model of computational anatomy [24, 25, 26, 27] in which the space of dense anatomies *I* ∈ ℐ is modelled as an orbit of a 3D template under the group of diffeomorphisms. Projective LDDMM models the sparse 2D histological observables not as an element of the orbit ℐ but rather a random deformation in dense 3D coordinates composed with a measurement projection to sparse coordinates. While LDDMM provides the geodesic metric [28, 29] on the orbit of 3D anatomies, there is no symmetry between the observable and the template, in general, and there should not be. This departs significantly from the symmetric methods [30, 31].

Alignment specifically of the modes of histology to MRI warrants two extensions of the basic model of Projective LDDMM. First, cross-modality similarity modelling is essential. Several strategies for representing image similarity have emerged including cross-correlation [32], mutual information [33], and local textural characteristics [34]. Our approach is to extend previous work [35, 36] by modelling a photometric transformation of histology to MRI with Mallat’s Scattering Transform [37, 38] to represent local radiomic textures at histological scales. Second, histological images carry large numbers of imperfections with tears, image stitching, and lighting variations. Extending previous work [15, 36], we introduce Gaussian mixtures models in the image plane of each histological slice to interpret image locations as matching tissue, background, or artifact. We proceed by way of the Expectation-Maximization (EM) algorithm [39] in estimating deformations that prioritize image matching at locations that are, in turn, estimated more likely to be matching tissue.

Here, we use Projective LDDMM to reconstruct the 3D geometries of two sets of 2D histological sections taken from the medial temporal lobe (MTL) of advanced cases of AD. As tau has exhibited stronger predisposition over A*β* for segregating to particular brain regions (ERC, CA1, subiculum) and layers (superficial) of cortex in AD [5], we use machine-learning based methods to detect and quantify neurofibrillary tangles (NFTs) from histological images. Modeling this data in a measure theoretic framework amenable to quantifying trends at different scales [40], we transport these detections to the 3D space via the correspondences yielded by Projective LDDMM.

## 1 Results

### 1.1 Projective LDDMM

In the random orbit model of computational anatomy [24], the hidden space of human anatomical images is modelled *I* : *R*^3^ → *R*^*r*^ as an orbit under diffeomorphisms of a template

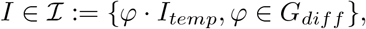

*G*_*diff*_ the group of diffeomorphisms *φ* : *R*^3^ → *R*^3^. The observables *J* : *R*^3^ → *R*^*q*^ are modelled as a random field with mean due to the randomness of diffeomorphic deformation and measurement process. For different problems of interest, the atlas image is ℝ^*r*^-valued with, for instance, *r* = 1 corresponding to single contrast MRI or *r* = 6 for diffusion tensor images (DTI) [41]. Likewise, observables are ℝ^*q*^ -valued with *q* = 3 for traditional histological stains or *q* >> 6 for alternative representations such as the scattering transform [37] encoding the meso-scale radiomic textures in histology images (see Section 2.2). In general, the range space of 3D templates versus targets do not have the same dimension, so *q* ≠ *r*.

Projective LDDMM is characterized by the fact that the observable is not dense in the 3D metric of the brain. Rather, the observable(s) result from either optical or physical sectioning, as in histological slice preparation, taking LDDMM into the projective setting akin to classical tomography [42, 43]. The sample measured observables *J*_*n*_(·), *n* = 1, 2, …are a series of projections *P*_*n*_ of *I*(·) on the source space *X* ⊂ *R*^3^ to measurement space *Y* ⊂ *R*^2^ (or *Z* ⊂ *R*^1^) defined through the class of point-spread functions associating source to target:

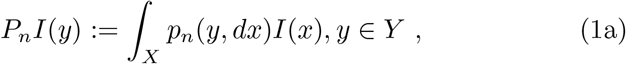

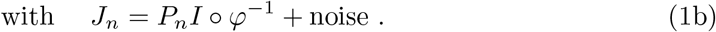

We adopt measure theoretic notation, *p*_*n*_(*y, dx*) for describing point-spreads to accommodate those taking the form of generalized functions, such as the delta dirac. Density notation *δ*(*x* − *x*_0_)*dx* corresponds to the measure notation 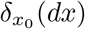, each evaluated against a test function *f* (*x*) ∈ *C*^0^, yielding *f* (*x*_0_).

The diffeomorphism *φ* is generated as the solution to the flow

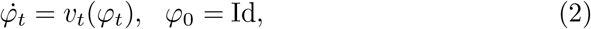

with velocity field *v*_*t*_, *t* ∈[0, 1] controlling the flow constrained to be an element of a smooth reproducing kernel Hilbert space (RKHS) 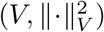 with the entire path square integrable 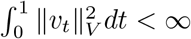 ensuring smoothness and existence of the inverse [44].

This gives us the first variational problem of Projective LDDMM.

#### Variational Problem 1

(Projective LDDMM)

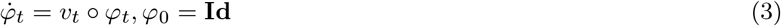

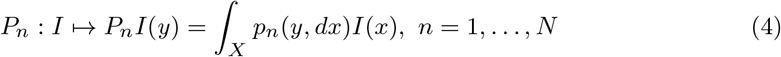

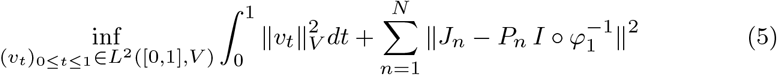

The model specific to our histology images projects the volumes *I*(·) on *X* ⊂ *R*^3^ to parallel sections *J*_*n*_(·) on *Y* ⊂ *R*^2^, along the third (*z*) dimension, with coordinates, *z*_*n*_, *n* = 1, · · ·, *N*. For this, we define ‘Dirac’ point-spreads from *δ*_*x*_ applying to infinitesimal volumes in space (*dx*) with *δ*_*x*_(*dx*) equal to 1 if *x* ∈ *dx*, and 0, otherwise. Our Dirac point-spreads 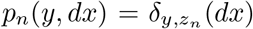 with (*y, z*_*n*_) = (*y*^(1)^, *y*^(2)^, *z*_*n*_) ∈ *R*^3^ concentrate on the planes:

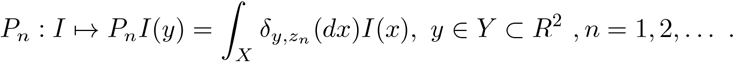

### 1.2 Histological Sectioning: Crossing Modality and Projective Plane Distortions

Because of tissue section preparation, we attach two transformations on the target planes. Deformation of the tissue section geometry associates diffeomorphisms to the histology, *ϕ*_*n*_ ∈ Φ : *R*^2^ → *R*^2^ both rigid and high-dimensional, expanding the dimensions first used in [14] for block sectioning.

For crossing from the range space of histology contrast to MRI we introduce a complete expansion, reparameterizing the histology with 48 dimensions via Mallat’s Scattering Transform [37, 38], and solve for the optimal linear predictor 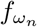, matching the scattering transform output to the scalar MRI. Via a *subsampled* Scattering Transform, color histology images *J*_*n*_ : *Y* → *R*^3^, *Y* ⊂ *R*^2^ are resampled at the resolution of MRI to 48-valued vector fields through alternating wavelet convolutions and nonlinear modulus operators across scales (see Appendix C):

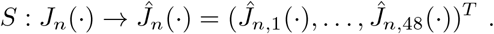

The linear predictor introduces the parametric class of contrast variations (*f*_*ω*_, *ω*∈ Ω) : *R*^48^ → *R* via polynomials with coefficients, *ω* (e.g. *ω*∈ *R*^49^), defined on the scattering output (e.g. 48-valued vector field) and constant offset. This gives Variational Problem 2.

#### Variational Problem 2

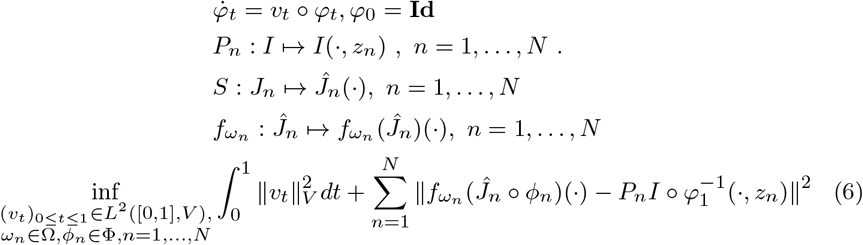

For non-rigid modeling of *ϕ*_*n*_, a penalty term 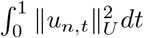 is added to (6), with 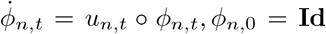. For estimating the cross-modality dimensions *ω*_*n*_, we have not mapped a large enough empirical sample to build an informative prior *π*_*ω*_ on those high dimensions. Therefore, we treat them in a maximum-likelihood setting, optimizing them with initial conditions of deformation and image plane dimensions fixed, then solve the variational problem over all of the other dimensions with the *ω*_*n*_ estimates fixed (see Section 2.2). This avoids collapse of the variational problem in these high dimensional settings.

### 1.3 Optical Sectioning, PET, and Parallel Beam Tomography: Ideal and Non-Ideal Planar and Linear Projections

Confocal optical sectioning reconstruct volumes *X* ⊂ *R*^3^ with models that are fundamentally 3-D point-spreads 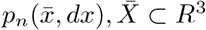 [45, 46], with imaging focused to *n* = 1, …, *N* measurement planes with significant blur out of plane. The mean field of the measurement volume 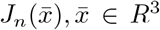 are given by the projections 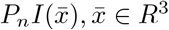 of (4).

Two-dimensional (2D) positron emission tomography (PET) introduces point-spreads *p*_*n*_(*y, dx*) for reconstruction which are less idealized supported over planes *Y* ⊂ *R*^2^ with uncertainty perpendicular to the line of flight but as well a second measurement the time-of-flight of the annihilating protons to the detectors [47]. Generally the point-spreads are modelled as cigar shaped two-dimensional Gaussians in the plane oriented by *N* angles *θ*_*n*_, *n* = 1, …, *N*, with high fidelity systems having *N* > 96, with standard deviation of uncertainty significantly larger along the lines of flight than perpendicular to them.

Classical parallel beam projection tomography reconstructs image planes *Y* ⊂ *R*^2^ via the Radon transform generated from sinograms indexed over a single space dimension dimension *Z* ⊂ *R*^1^ arising from idealized line integrals [48]. Define the set of oriented lines in *R*^2^ parametetrized by their angles (*θ*) and offsets from the origin (*z*), *L*_*θ*_(*z*) = {(*y*^(1)^, *y*^(2)^)} ⊂ *R*^2^ with

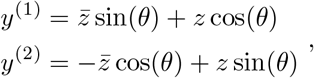

for 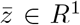. The mean field of the measurements *J*_*n*_(*z*), *z* ∈ *R*^1^ are given by the line integrals 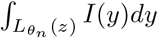. To satisfy the basic sampling theorems for tomographic reconstruction [49], *N* sampling angles *θ*_*n*_, *n* = 1, …, *N*, akin to N slices in histology, are selected determined by the resolution required for the reconstruction. The point-spreads corresponding to the line integrals becomes 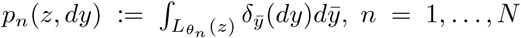, with the projections *P*_*n*_*I*(*z*), *z* ∈ *R*^1^ of (4), (see Appendix A).

### 1.4 Biological Results: Multi-scale Maps of Tau Tangles in 3D

We present, in this section, the geometric and neuropathological results for corresponding pairs of histology images and MRI, taken postmortem, of MTL tissue from two cases of advanced AD. To acquire these data, several steps are required (see Section 2). Solutions to Variational Problem 2 (Equation 6) yield geometric reconstruction of histologically stained tissue in 3D. Figure 1 illustrates the two sets of 35 individual digitized sections on which NFTs were detected and geometric mappings to 3D were estimated. Subsequent coordination of both MRI and mapped histology with the Mai Paxinos Atlas is demonstrated in Figure 2, with coronal Mai views shown for an example intersecting histological slice.

**Fig. 1.**
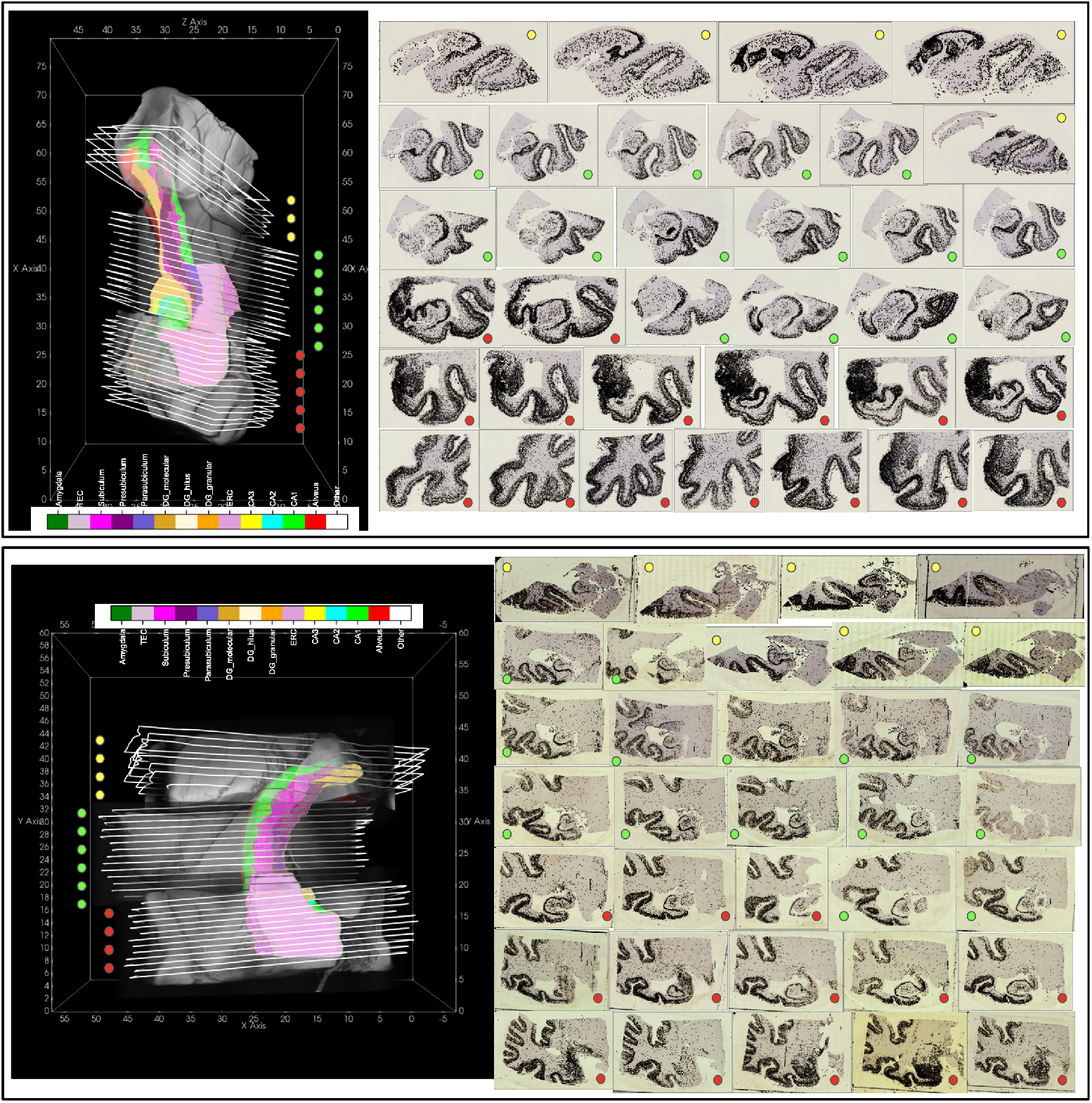
Complete datasets of PHF-1 stained histology sections for 3 blocks of MRI for each brain sample. 3D MRI shown with manual segmentations of MTL subregions (left). Boundary of each histological section on right sketched in position following transformation to 3D space (left). Detected tau tangles plotted over each histology slice (right).

**Fig. 2.**
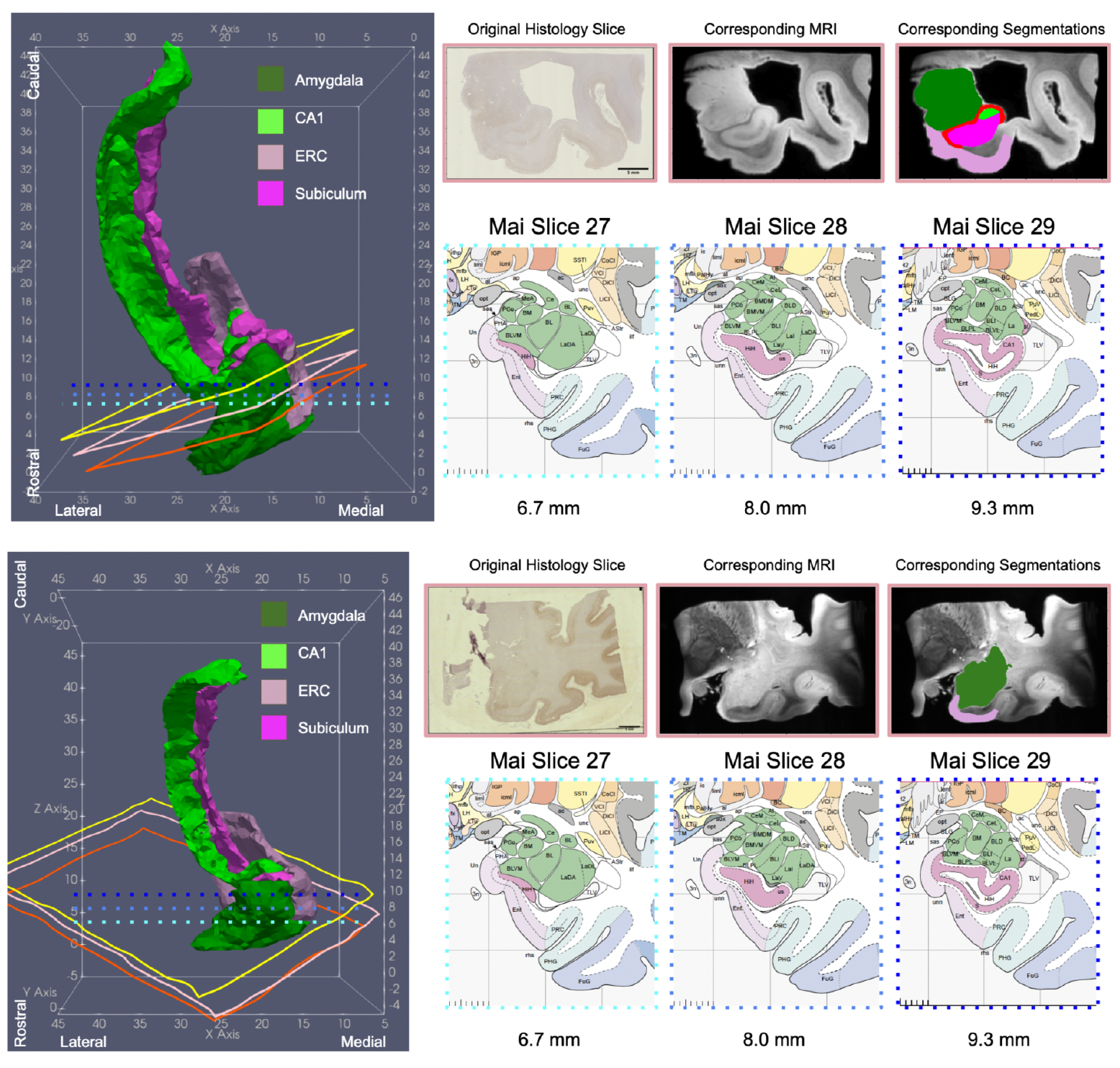
3D Reconstruction (left) of 4 MTL subregions for two advanced AD brains in the coordinate space of the Mai Paxinos Atlas. Corresponding section of histology and MRI (top right) shown and intersecting coronal planes taken from the pages of the Mai Atlas (bottom right).

Figure 3 illustrates 4 representative samples of manual segmentations compared between histology and MRI deformed to 2D. The protocol for segmentation is described in section 2.5. Alignment accuracy was measured from these comparisons with Dice overlap and 95th percentile Hausdorff distance for MTL subregions of interest (see Appendix B). The latter measure ranged from 1.0 mm to 2.0 mm across amygdala, ERC, CA1, and subiculum.

**Fig. 3.**
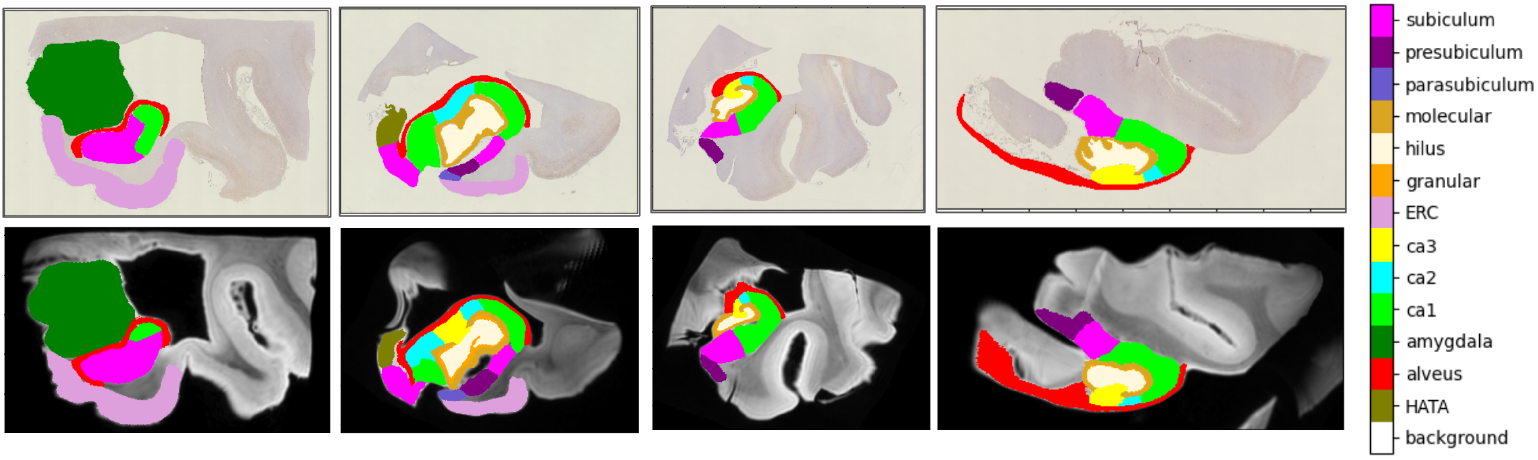
Selected histology slices with 2D segmentations (top row) ordered left to right as rostral to caudal. Corresponding MRI slices with 3D segmentations mapped to 2D via transformations *φ, ϕ*_*n*_ (bottom row).

Accuracy of tau tangle detections was evaluated both at intermediate and final steps of our detection algorithm. Estimates of accuracy in per pixel tau probabilities were computed using 10-fold cross validation on the entire training dataset for each brain sample. Table F3 (see Appendix F) shows accuracy metrics for one brain sample, with mean AUC and of 0.9860 and accuracy of 0.9729.

Accuracy of individual counts of NFTs as output following segmentation by the watershed algorithm (see Section 2.6) were estimated by comparison to manually annotated patches of ERC tissue reserved for validation. Between 5 and 20 mm^2^ patches in the region of the ERC on 10 roughly consecutive slices of one brain were selected for annotation. Per pixel annotations were completed by a single individual over the course of 2-3 weeks. Each patch totaled approximately 2, 500, 000 pixels, yielding a total of 25 million for the 10 patches–on the order of the number of voxels for 20 whole brain MRIs at 1 mm resolution. Densities of NFTs (counts per cross-sectional tissue area) for each patch were computed using counts of manually annotated tangles vs. algorithmic output and rescaled to the range, [0, 1], for accurate comparison. Slice-by-slice relative densities are illustrated in Figure 4, exhibiting high levels of similarity between human and machine generated NFT densities.

**Fig. 4.**
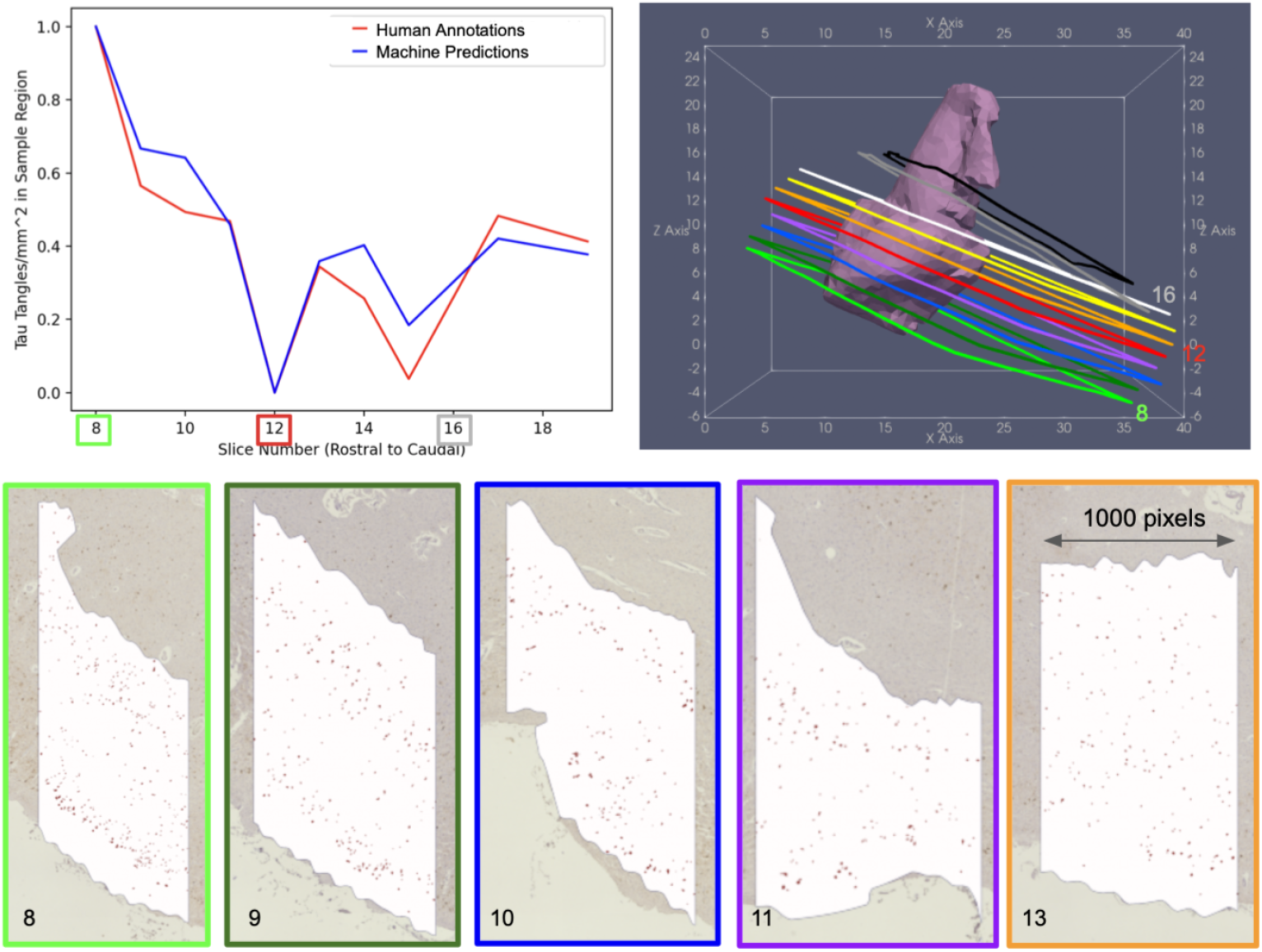
Tau tangle densities within patches of ERC computed from manual annotations (red) and machine prediction (blue), normalized to [0, 1] in each case (left). Outlined sections of histology from which validation set of ERC patches was taken. Sections plotted post transformation in coordinate space of Mai atlas with 3D reconstruction of total ERC (right). Example patches in ERC (white) with NFTs annotated (red) for five slices (bottom).

Counts of detected NFTs, cross-sectional tissue area, and MTL subregion (from MRI deformed to 2D) were computed in the space of histology slices. Average NFT densities per MTL subregion, tallied from all histology slices per brain sample, showed highest amounts of NFTs in amygdala, ERC, CA1, and subiculum for both advanced AD samples (see Figure 5,7).

**Fig. 5.**
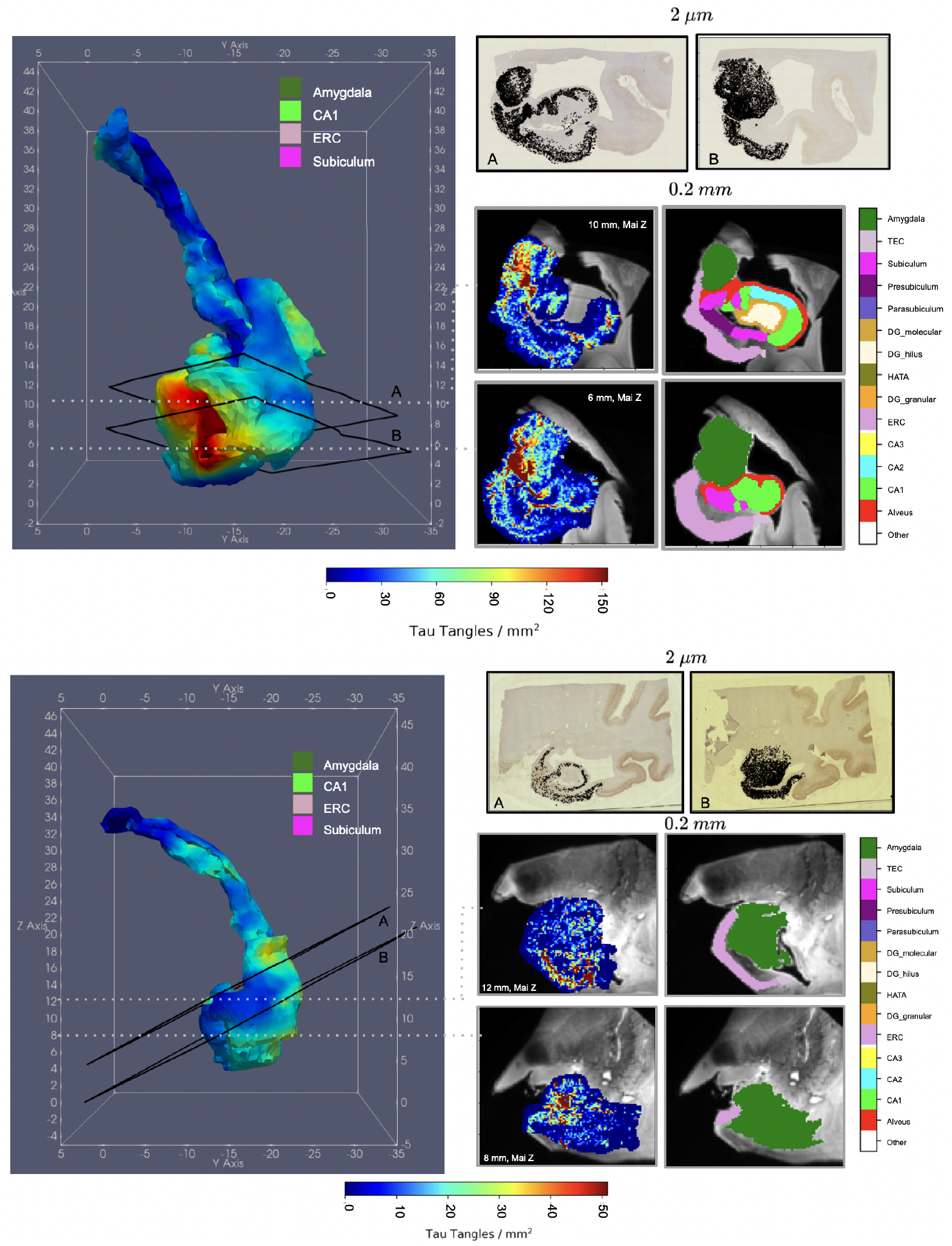
Reconstructed NFT densities in 3D MTL of advanced case of AD. Densities within subset of MTL (amygdala, ERC, CA1, and subiculum) computed over surface of each structure (left) and within the dense metric of the 3D MRI (right). MRI slices correspond to coronal slices in Mai Paxinos atlas.

Histological data was modeled as discrete particle measures and subsequently transported to the 3D space of the Mai Paxinos atlas via estimated transformations, *ϕ*_*n*_, *φ* (see Section 2.7). Diverse modes of resampling yielded distributions of NFT density within, over, and between MTL subregions (see Appendix G). Resampling via Gaussian kernels yielded smoothed NFT densities computed within the dense metric of the brain at approximate resolutions of MRI. Resampling via nearest neighbor kernels projected particle measures to the surface of segmented MTL subregions, such as amygdala, ERC, CA1, and subiculum for visualization in 3D. These are both shown in Figure 5.

Average NFT densites per region are summarized in Figure 7. Tau pathology localized not just *to* particular regions (e.g. amygdala, ERC, CA1, subiculum), but within them. As illustrated in Figure 6, high densities of NFTs in amygdala and ERC concentrated particularly at the border between the two structures, with tau migrating to the inferior, medial boundary of the amygdala. NFT density also appeared to segregate within the hippocampus with highest densities achieved in the anterior third of the hippocampus in both brain samples (see Figure 7).

**Fig. 6.**
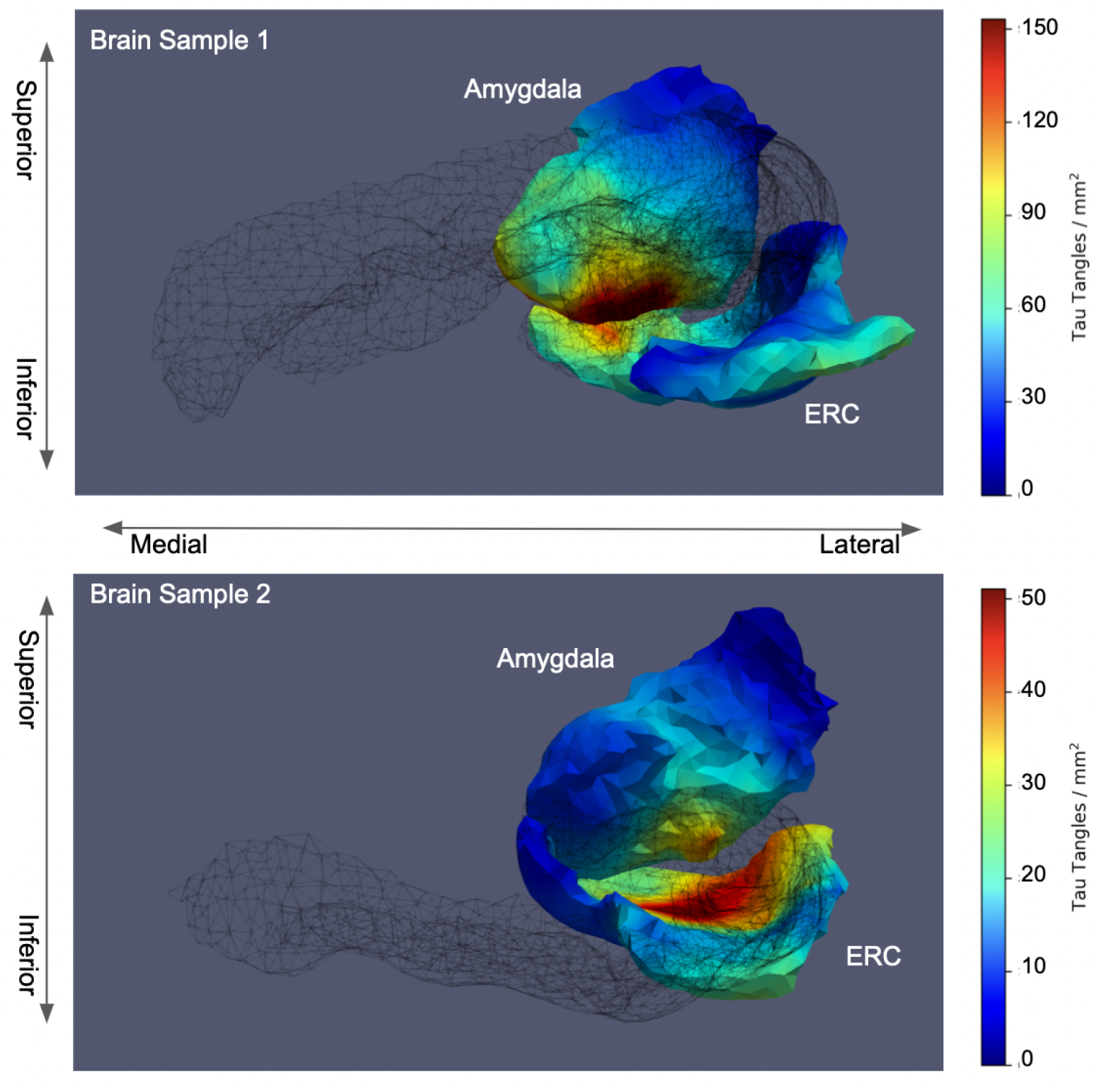
Posterior view of amygdala-ERC boundary in brain sample 1 (top) and brain sample 2 (bottom). NFT densities projected to and smoothed over surface of each structure independently. Outline of CA1 and subiculum surfaces shown in black mesh.

**Fig. 7.**
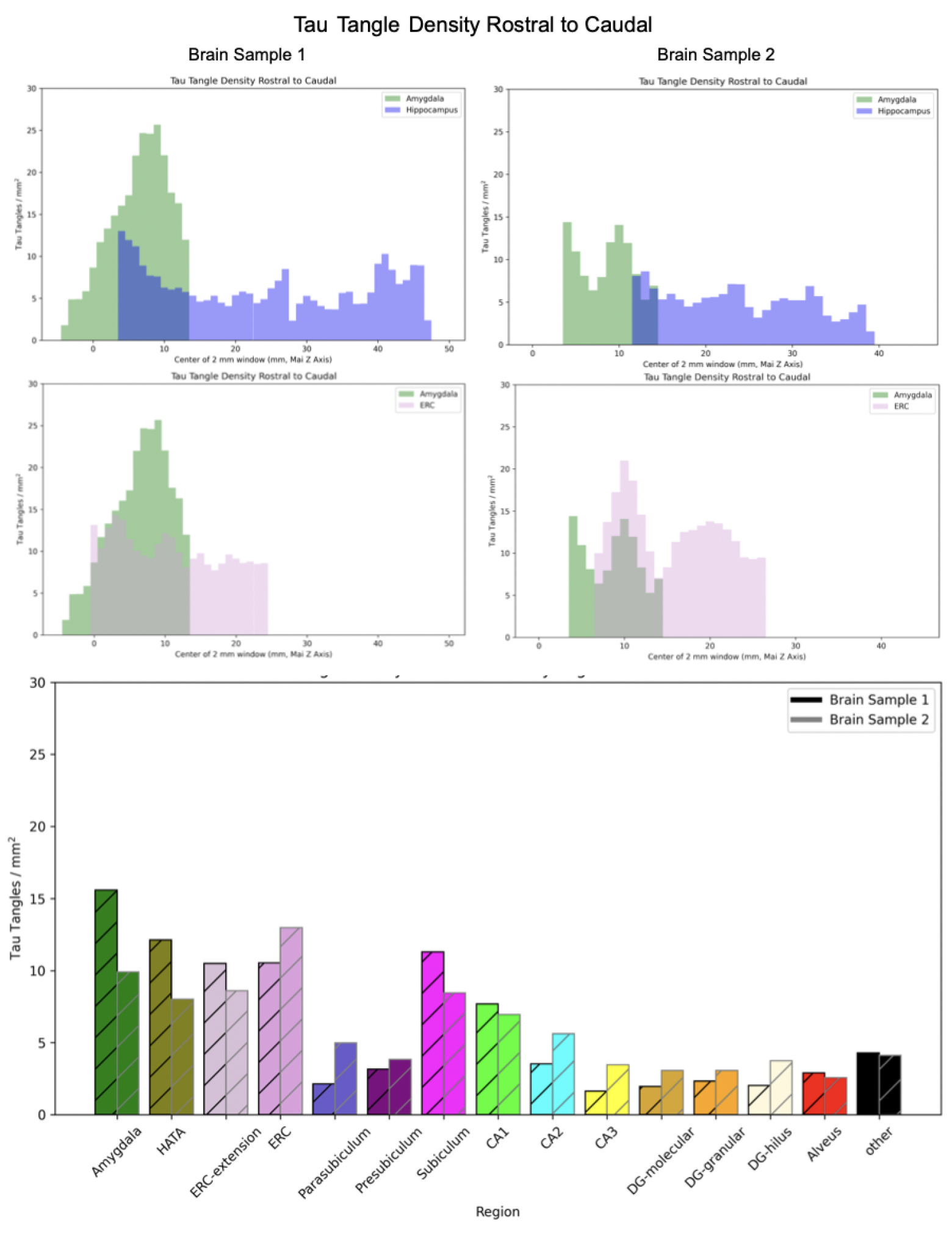
NFT densities along Mai Z axis (rostral to caudal) in two cases of advanced AD. First two rows illustrate NFT densities within 2 mm sliding window along Mai Z axis for amygdala and whole hippocampus (top) and amygdala and ERC (middle). Bottom row illustrates global average NFT density within each MTL subregion. All densities calibrated against simultaneously stained slices from both brains (see Section 2.6).

## 2 Methods

## 2.1 Algorithm for Solving Projective LDDMM with In Plane Transformation

To solve the Variational Problem 2 for 3D atlas, *I*, and set of 2D targets (*J*_*n*_, for *n* = 1, …, *N*) we formulate an algorithm that alternately optimizes for the deformation in 3D space and the geometric transformation in 2D space, while holding the other fixed. The algorithm can be implemented to incorporate increasing complexity as needed first for crossing modalities and second for crossing resolutions, as needed, for instance, to map 3D MRI to 2D histological slices, as is presented here. In its simplest form, *I* and *J* are of the same modality, yielding 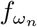 fixed as identity, and *ϕ*_*n*_ is modeled as a rigid motion in plane, as in Lee et. al mapping histological sections to an atlas of the mouse brain [14]. Transformations *φ, ϕ*_*n*_ for *n* = 1,…, *N* are estimated following Algorithm 1.

### Algorithm 1

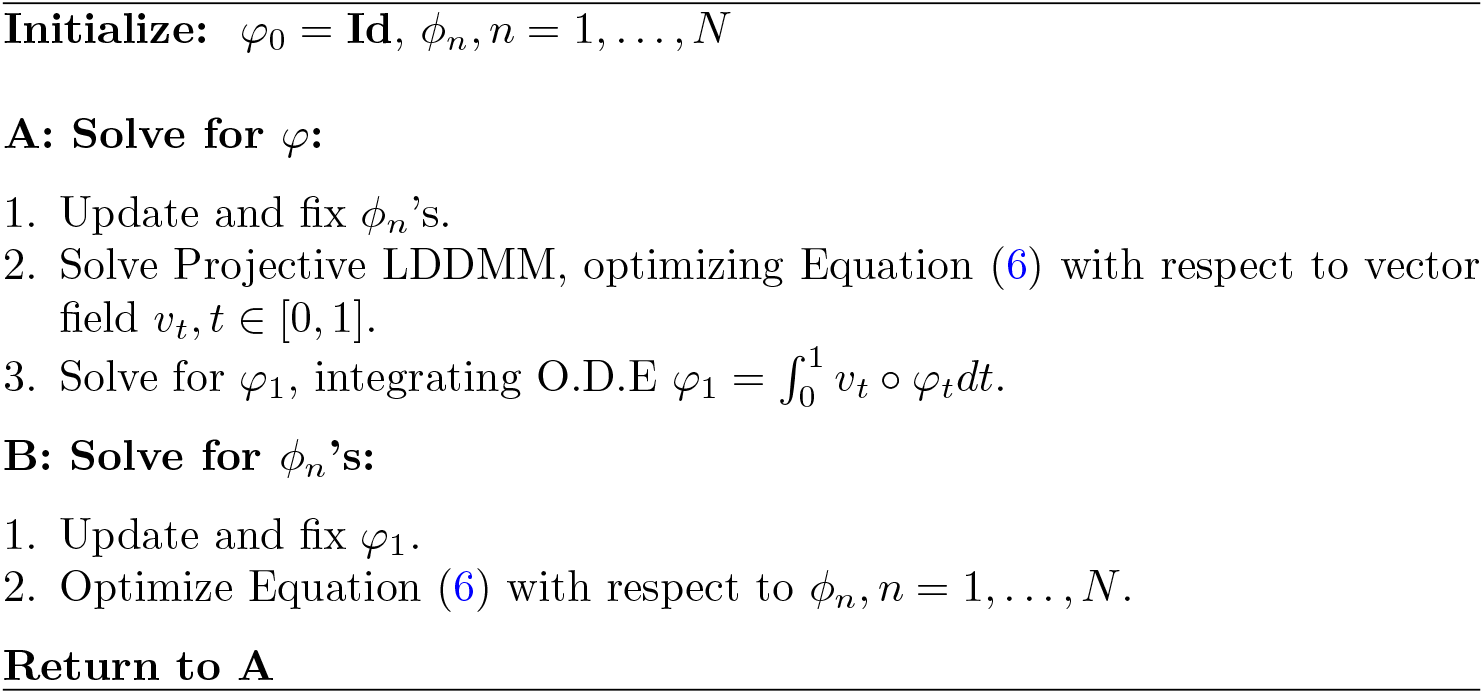

When *ϕ*_*n*_’s are broadened to non-rigid diffeomorphisms, as in [15], each *ϕ*_*n*_ is estimated in step B via a separate iteration of LDDMM for each target *J*_*n*_. Here, *ϕ*_*n*_’s encapsulate both rigid and non-rigid components. Separate gradient based methods are used to update each component in step B with velocity fields updated using Hilbert gradient descent as in [50] and linear transform parameters updated by Gauss-Newton [51].

### 2.2 Optimization Algorithm for Solving Projective LDDMM Crossing Modalities and Resolutions via Scattering Transform

Crossing modalities at similar resolution (e.g. 3D MRI and downsampled 2D histology slices) requires re-introduction of 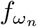 as a polynomial mapping between range spaces of template and target, giving a similar formulation to that used in Tward et. al [15]. In this work, we introduce the Scattering Transform [37] for crossing modalities at *differing* resolution. We model contrast variations between histology and MRI by (i) representing local radiomic textures via 48 dimensions of histological scales using Mallat’s Scattering Transform [37, 38], (ii) dimension reduction projecting the 48 dimensional scattering transform onto a 6-dimensional PCA basis, and (iii) solving for the optimal linear predictor 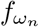 matching the scattering transform 6-dimensions to the scalar MRI.

Histology images *J*_*n*_ : *Y* → *R*^3^, *Y* ⊂ *R*^2^ are color images resampled to 48-valued vector fields via the Scattering Transform (see Appendix C): 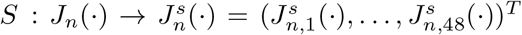 defined at the resolution of MRI. A basis, *b*_1_,…, *b*_48_, is computed offline for the space of Scattering coefficients using PCA. A 6-valued vector image is generated from 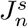 as the projection onto the 6 largest eigenvalue basis elements, yielding the predictor, 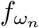 as affine including a constant offset:

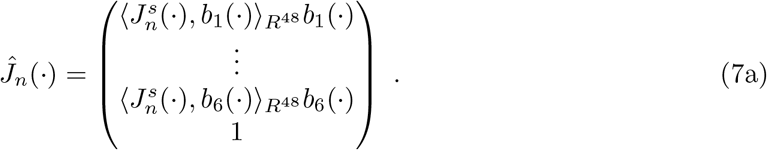

The mean field histology *f*_*ω*_ parameterized by linear weights *ω* ∈ *R*^7^ becomes:

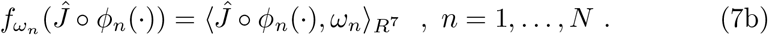

Figure 8 shows a mean field section using the scattering transform. These linear weights *ω*_*n*_ are estimated from initialized *ϕ*_*n*_’s and *φ* following Algorithm 2. Initializations of *ϕ*_*n*_ and *φ* are estimated following the approach in Tward et. al [36] in which cubic polynomials are used to match MRI range space to histology range space. Solutions for *ω*_*n*_ are then obtained using the pseudo-inverse:

**Fig. 8.**
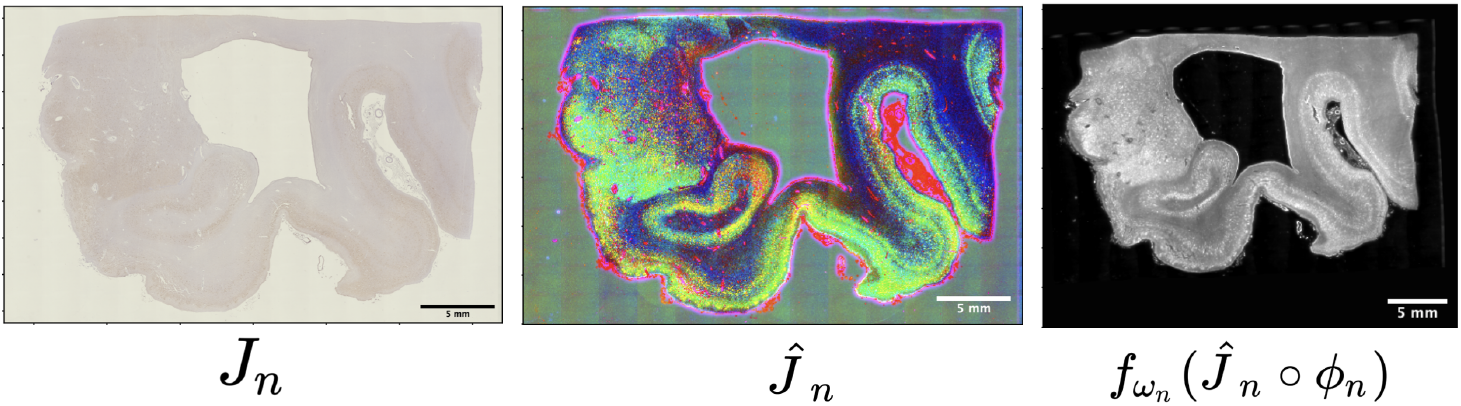
Initial histology image *J*_*n*_ at 2 *μ*m resolution (left). First three coefficents of scattering image 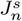 projected to 6-dimensional PCA basis (center). Output of linear predictor mapping deformed image *Ĵ*_*n*_ ∘ *ϕ*_*n*_ to grayscale MRI intensity range (right).

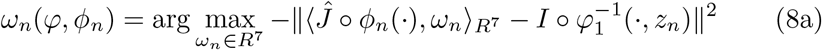

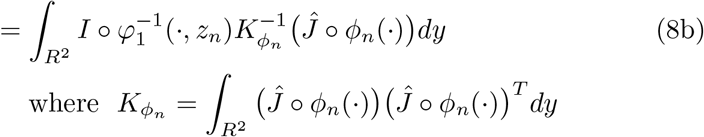

#### Algorithm 2

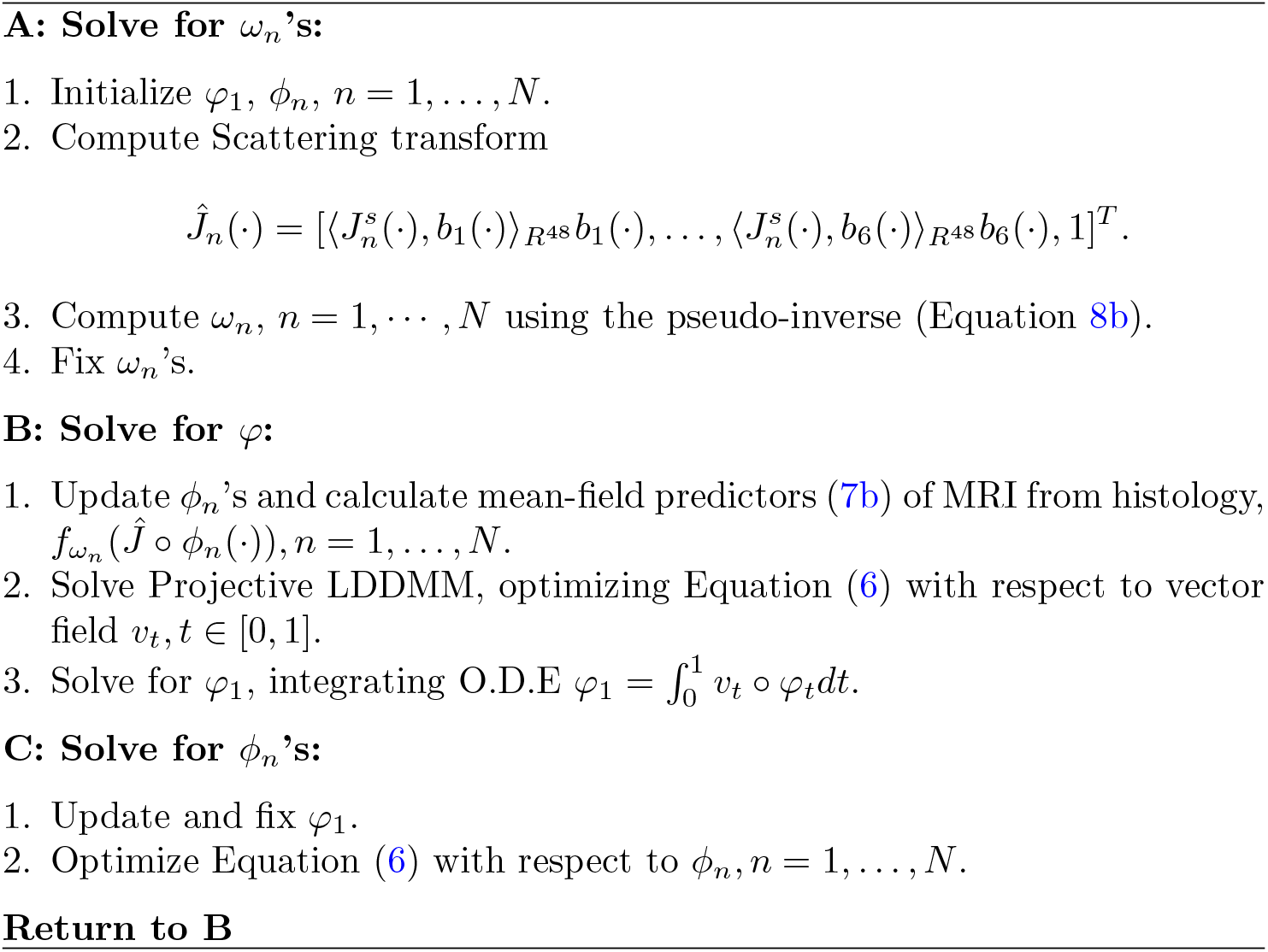

### 2.3 Algorithm for Solving Projective LDDMM With Distortions and Missing data via EM Algorithm

The last element of complexity we introduce aims to account for the many distortions and subsampled missing data sections not wholly accounted for by geometric transformations, *ϕ*_*n*_. We formulate a weighted, least-squares problem by introducing weights representing different model interpretations of the pixels in each histology tissue section: measurements (e.g. tissue foreground), artifacts (e.g. tears and distortion), or background, following the example in Tward et. al [36]. Each model is characterized as a Gaussian with standard deviation, *σ*_*M*_, *σ*_*A*_, *σ*_*B*_ and mean:

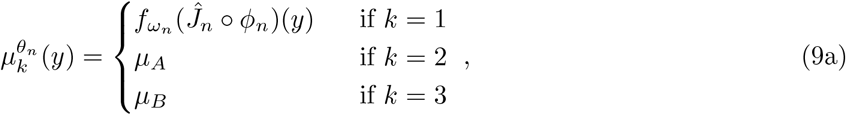

with *μ*_*A*_ and *μ*_*B*_ representing artifacts and background. Weighted least-squares interprets the images weighing each model 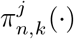, with 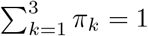, giving

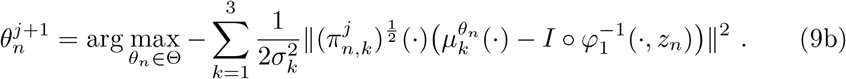

The weights arise from the E-step of an Expectation-Maximization (EM) algorithm [39], that we use for estimation of parameters *θ*_*n*_ = *ϕ*_*n*_ in step C of our Algorithm 2. Weights are a function of the previous parameters 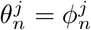 hence giving the iteration: selecting at each point in the image the appropriate model for giving the spatial field of weights. This iteration corresponds to a Generalized EM (GEM) algorithm [39] (see Appendix D for proof). The results highlighted in Section 1.4 were generated following the approach of this section. For select slices in each brain sample, in plane *ϕ*_*n*_’s were estimated in step C of Algorithm 2 as spline transformations via manual landmark placement.

### 2.4 Specimen Preparation and Imaging

Brain tissue samples were prepared by the Johns Hopkins Brain Resource Center. The demographics and pathological staging of each sample analyzed are summarized in Table E, Appendix E. From each formalin immersion fixed brain, a portion of the MTL including entorhinal cortex, amygdala, and hippocampus, was excised in 3-4 contiguous blocks of tissue, sized 20-30 mm in height and width, and 15 mm rostral-caudal (see reconstructed MRI of tissue blocks, Figure 1).

Each block was imaged with an 11T MR scanner at 0.125 mm isotropic resolution and then cut into two or three sets of 10 micron thick sections, spaced 1 mm apart. Each block yielded between 7 and 15 sections per set. Sets of sections were stained with PHF-1 for tau tangle detection, 6E10 for A*β* plaque detection, or Nissl, and digitized at 2 micron resolution.

### 2.5 Segmentations of MTL Subregions

In all brains, MTL subregions were manually delineated using Seg3D [52]. Individual block MRIs were rigidly aligned using an in-house manual alignment tool, and per voxel labels were saved for the composite MRI for each brain. Delineations were deduced from patterns of intensity differences, combined with previously published MR segmentations [53, 54, 55] and expert knowledge on the anatomy of the MTL. The established borders were applied in three other brains, showing consistent results (in preparation, EX, CC, SM, DT, JT, Alesha Seifert, Tilak Ratnanather, MA, MW, and MM). In one brain sample, corresponding regional delineations were drawn on all histology sections stained with PHF-1 (see Figure 3). Delineations were based on visible anatomical markers and were afterwards confirmed with a corresponding Nissl-stained set of sections. In each of these sections, cytoarchitectonic borders between areas of interest were indicated, independently from the other datasets, using an previously published cytoarchitectonic accounts of the MTL [56, 57, 58, 59, 60, 61]. Labels were assigned per pixel to 4x-downsampled histology images at a resolution of 32 microns and used to evaluate accuracy of registration (see Section 1.4). Regions of interest include amygdala, entorhinal cortex (ERC), cornu ammonis fields (CA1, CA2, CA3), and subiculum (see Figure 1).

### 2.6 Density Determination of NFTs

Patterns of tau pathology are summarized as total counts of tau tangles (NFTs) per mm^2^ of cross-sectioned tissue. NFT counts were computed using a 2-step algorithm: (1) prediction of per pixel probabilities of tau and (2) segmentation of these probability maps into discrete NFTs.

As described previously [62, 36], we used a convolutional neural network to model and predict probabilities of being part of a tau tangle for each pixel in a digital histology image. To capture larger contextual features as well as local information for producing per pixel probabilities at high resolutions, we trained UNETs [63] with the architecture described in Table F2 (see Appendix F). Given differences in staining intensity between brain samples, we trained separate UNETs, each with the same architecture, for each brain sample. Training data per brain was generated on every third slice of histology. Between 8 and 24 sample zones, sized 200-by-200 pixels were selected at random until 8 zones covered tissue (not background). Every pixel in each zone was manually annotated, 1 or 0, as part of a tau tangle or not.

Counts of NFTs in each histology slice were generated by segmenting the probability maps output from the trained UNET. Segmentations were computed using an opensource implementation of the watershed algorithm [64] to extract connected components with “high probability” of tau. Each component was defined as an individual NFT, with center, area, and roundness computed as features.

In the results above (Figures 4,5,6), NFT densities are reported on different scales for each brain sample as differences in staining intensity, timing, and handling of tissue samples for each brain resulted in different scales of absolute counts of NFTs. To compare NFT densities across brain samples, subsets of 4-5 additional sections from each brain were selected in the rostral hippocampus at approximate locations of original sections. These two subsets were stained simultaneously to achieve consistency between them. A UNET was trained on the original training data from both brains and NFTs were detected and summed across each of original and new sets of histology slices. Ratios of tau tangles detected on the original vs. new version of each section were computed, yielding average ratios of 4.4 and 1.7 for brain samples from subjects 1 and 2, respectively. Finally, assuming consistent overestimation in tau tangle density measures across sets of slices, measures along the rostral-caudal axis and in MTL subregions were rescaled in each brain according to the average ratio factor to produce comparable NFT densities (see Figure 7).

### 2.7 Particle Representation of Histological Data

We model histology data at the microscopic scale following the generalized measure approach in [40] where each particle of tissue carries a weighted Dirac measure over histology image space and a Dirac measure over the feature space 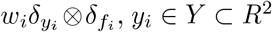 and ℱ = *R*^2ℓ^. Weights reflect sampled tissue area captured in each particle measure, defined at the finest scale (*μ*^0^) as cross-sectional area in the histology plane *w*_*i*_ ∈ {2*μm*^2^, 0}, computed with thresholding using Otsu’s method [65]. The first ℓ dimensions of *f*_*i*_ ∈ ℱ denote the number of tau tangles in each of ℓ MTL subregions. The second ℓ dimensions denote the fraction of sampled tissue area (*w*_*i*_) within each of ℓ MTL subregions. At the finest scale,

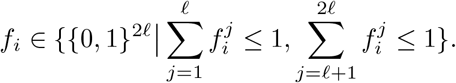

We transfer measures via diffeomorphisms *ϕ*_*n*_ and *φ* and rigid transformation to the space of template *I* and the Mai Paxinos Atlas. Discrete weights *w*_*i*_ adjust according to in plane expansion/contraction of cross-sectional tissue area, with adjustment at the fine scale by *ϕ*_*n*_ given by the varifold action:

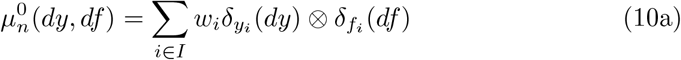

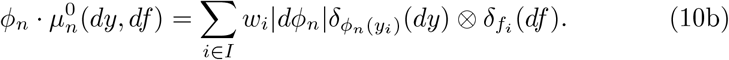

To cross scales we use the fundamental decomposition of the particle measures

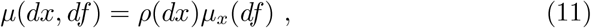

with *ρ* being the density of the model and *μ*_*x*_ the field of conditional probabilities on the features. Our tranformation across scales non-linearly rescales space and smooths the empirical feature distributions on the features. Spatial resampling is determined by *π*(*x, x*′), the fraction that the particle at *x* assigns to the particle at *x*′, with 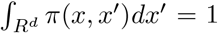. The smoothing on the field of conditional probabilities occurs according to choice of kernel *k*((*y, α*_*y*_), ·):

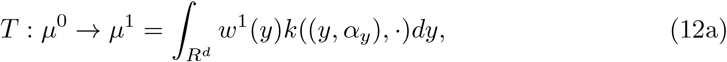

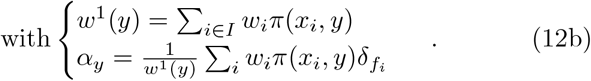

Spatial resamplings at MRI resolutions (0.125 mm) and over surface boundaries of MTL subregions were achieved through isotropic Gaussian resampling and nearest neighbor resampling, respectively, through choice of *π* (see Appendix G). Feature reduction occurs via maps *β*↦ *ψ*(*β*) ∈ ℱ′, for probability measures *β* with 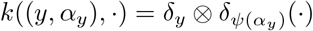 giving

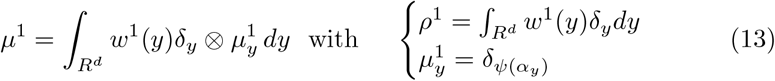

Here, *ψ*(·) reduces feature dimension by taking empirical distributions *α*_*y*_ over each of 2ℓ dimensions to expected first moments for each corresponding dimension, giving ℱ′ = *R*^2ℓ^ with:

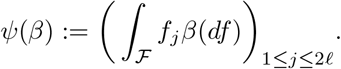

Total NFT density is computed from the sum of the first ℓ features while NFT density per region is computed from the ratio of feature value *j* to ℓ + *j* for any of *j* = 1, · · ·, ℓ MTL subregions.

### 2.8 Surface Smoothing using Laplace Beltrami Operator Basis

Spatial variations in NFT density within MTL subregions are visualized as smooth functions over the surface of each corresponding region. Particle mass belonging to a given subregion volume is “projected” to the surface boundary using a nearest neighbor kernel for *π*, as defined in Equation 12b (see Section 2.7):

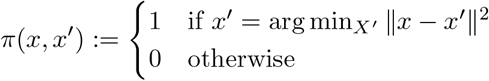

We construct functions, *g*_*τ*_ (*x*) and *g*_*a*_(*x*), to represent the total number of NFTs and cross-sectional area of tissue from discrete particle measures of particles projected to the surface vertices *x*_*i*_ ∈*V*. We generate smooth representations of NFT density 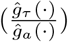 using Laplace-Beltrami on each surface [66]:

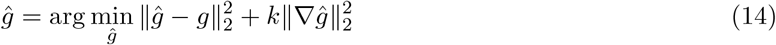

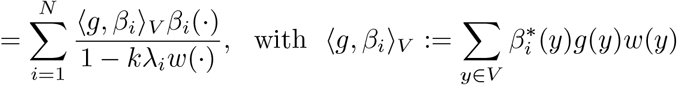

for both *g*_*τ*_ (·) and *g*_*a*_(·), where ℬ := {*β*_1_, …, *β*_*N*_} is a basis for the Laplace-Beltrami operator, and *k* the smoothing constant (see Appendix H).

## 3 Discussion

These findings demonstrate that 3D maps of tau tangle density at mm and micron resolution show a strong spatial predominance of tau in the rostral third of the MTL, with especially high densities measured in the amygdala. They can be considered a ‘proof of concept’ of an approach comprised of three key components. (1) The definition of Projective LDDMM is the key contribution of this work, as a class of image-based diffeomorphometry methods for reconstructing sparse and irregularly sampled data in the dense metric of the 3D brain. (2) The coupling of geometric and contrast transformations generalizes these methods further to images of different modalities. (3) The use of a measure theoretic framework to model data following reconstruction (i) preserves data mass and values and (ii) allows for resampling the dense metric to generate statistical distributions at arbitrary resolution. Evidence of sparsity in original data measurements is maintained in the preservation of data mass while reconstruction and resampling within the dense metric offers smooth interpolation between the data values.

We demonstrate the success of this approach at reconstructing spatial profiles of tau tangle density in two MTL samples from individuals with advanced AD. By computing NFT “density” as counts of tau tangles per cross-sectional area, we maintain data values reflective of the original 2D histological space in which they were measured. Reconstruction within the dense 3D metric of the brain allows for resampling of these measures at micron and mm resolution along both regular and irregular sampling “grids”, such as over surface boundaries of MTL structures. Reconstruction within the spatial coordinate system of a well-known atlas (e.g. Mai atlas) allows for consistent resampling across brain samples, such as along the rostral-caudal axis, for comparing profiles of pathology.

Both reconstructions illuminate AD as a spatially-oriented disease, which further motivates the need for generating such 3D spatial reconstructions to uncover these significant patterns. Specifically, two main patterns in tau pathology emerged for both brain samples: (1) high levels of NFT density in amygdala and ERC, and (2) a high-to-low gradient of NFT density along the rostral-caudal axis of the hippocampus (see Figure 7), localizing NFT density to the regions where it abuts the Amygdala and ERC. Though both samples reflect end-stage pathology, the rostral-caudal gradient is consistent with the spread of tau outlined in Braak staging [5, 6], and supports recent work by Yushkevich et. al that found similar segregation of tau pathology in the anterior vs. posterior portions of hippocampal structures in cases of earlier stage disease [35].

The role of the amygdala in AD has historically been under less investigation than the MTL subregions initially highlighted by Braak (e.g. TEC, ERC, CA fields, and Subiculum) [5]. Nevertheless, a number of studies have suggested its involvement not just with AD but other neurodegenerative diseases involving misfolded proteins [67, 68, 69]. Its emergence here as a key reservoir of tau pathology complements these studies as well as Yushkevich et. al, who found high burdens of tau pathology in the amygdala [35]. Furthermore, previous work from our group in MRI diffeomorphometry suggested a similar effect of AD pathology in the amygdala as in the ERC based on shape markers reflective of atrophy [70, 71]. Both basolateral and basomedial regions of the amygdala demonstrated significant atrophy, consistent with the appearance of higher tau densities, here, in the amygdalar regions adjacent to the ERC (Figure 6).

Though this work has been limited to the analysis of two brain samples, the methods we’ve described and the distributions of tau pathology that have emerged suggest a number of avenues for expanding on these efforts. Projective LDDMM, as we’ve formulated it here, can easily see value in applications to light sheet microscopy, spatial transcriptomics, and other emerging imaging modalities in which sparsely sampled data sets must be reconstructed in the dense space of some atlas for full appreciation of the spatiotemporal resolution of these modalities. Within the realm of AD research, we foresee its use in drawing additional correspondances between microscopic neuropathology and imaging modalities, such as MRI, PET and DTI, in the validation and development of biomarkers.

The segregation of tau pathology to the rostral most third of the hippocampus, the amygdala, and the ERC also suggests direction of future study in AD. First, MRI images that routinely capture the entirety of the hippocampus might be refined to target this smaller section of the hippocampus, amygdala and ERC, offering greater specificity to AD and sensitivity to neuropathological changes, as higher resolutions could be achieved with a narrower region of capture. Second, the spatiotemporal distribution of neuropathology at the earliest stages of AD remains evasive and yet most significant in efforts to achieve earlier diagnosis. Emerging evidence of the neuropsychiatric syndromic complex known as Mild Behavioral Impairment (MBI) amongst individuals prior to the onset of AD [72, 73] suggests a role for the amygdala, as an emotional and behavioral organ of the brain, particularly in these early stages. Hence, we are currently using our methods to reconstruct the distributions of tau pathology in earlier cases of AD to elucidate whether the amygdala shows similarly high levels of tau in these stages.

In summary, AD research, like so many other fields of neurodegeneration and neurobiology, harbors a disparity between knowledge, communication, and scientists working at the microscopic level and those at the macroscopic level. This disparity manifests prominently in incomplete understanding of the 3D spatiotemporal progression of pathology (tau and A*β*) in AD. This, in turn, has prohibited the progression of MRI biomarkers for earlier diagnosis of AD, as they have not been adequately linked to corresponding 3D patterns of neuropathology. We have developed the method we call Projective LDDMM to address this gap.

## Supplementary information

This manuscript is accompanied by a set of Supplementary Notes (Appendices) (referenced A-H in the main text).

## Acknowledgments

This research was funded in part by grants from the National Institutes of Health (U19-AG033655, P30-AG066507, P41-EB031771, R01-EB020062 (MM), T32-GM136577 (KS), U19-MH114821, R01-NS074980-10S1, RF1MH126732, RF1MH128875, RF1MH28888 (DT)), the Kavli Neuro-science Discovery Institute (MM,DT), and the Karen Toffler Charitable Trust (DT).

## Declarations

### Ethics Approval

This work was deemed not Human Subjects Research by the Johns Hopkins University IRB.

### Consent to Participate

Not applicable.

### Consent for Publication

Not applicable.

### Availability of Data and Materials

Due to size limitations, imaging data is available upon request.

### Code Availability

Code used for training and applying UNET in tau tangle detection can be found here: https://github.com/twardlab/ADproject. Code for solving Projective-LDDMM, building measure theoretic data representations, and resampling across scales can be found here: https://github.com/kstouff4/projective-lddmm.

### Authors’ Contributions

MA, SM, and JT recruited brain samples and oversaw preparation of histology and imaging. MW, CC, and EX manually segmented histology images and MRI. KS, MM, and DT developed methods. KS and DT implemented algorithm. KS and MM drafted manuscript. All authors contributed to editing final manuscript.

## Appendix A

### Classical Tomography

The Radon Transform is used in classical tomography [48] to describe the generation of sinograms as projected images at different angles. The transform is typically written in functional notation as an integral along one dimension, with Lebesgue measure as:

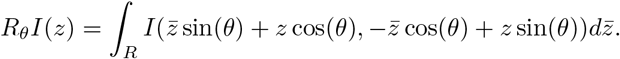

It can also be modeled as the integral over the line parametrized by two dimensions, an angle *θ* and affine offset *z* from the origin:

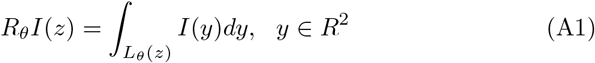

with the line defined by the paired angle and affine offset (*θ, z*) given explicitly by all *y* ∈ *R*^2^ such that:

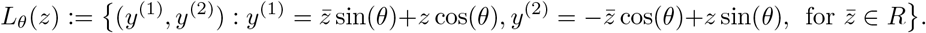

In the notation introduced in Section 1.1, we extend this to an integral over all of *R*^2^, using Dirac delta measures that assign nonzero measure only to lines in this same set:

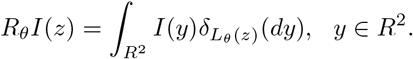

The line integral indexed by (*θ, z*) is modeled as a single projection *P*_*n*_*I*(*z*), *z ∈ R* indexed by a set of *n* = 1, …, *N* and is given as in, equation 4, with point spread, 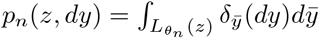, as defined in section 1.2. We show below its equivalence to the line integral as defined in above (A1).

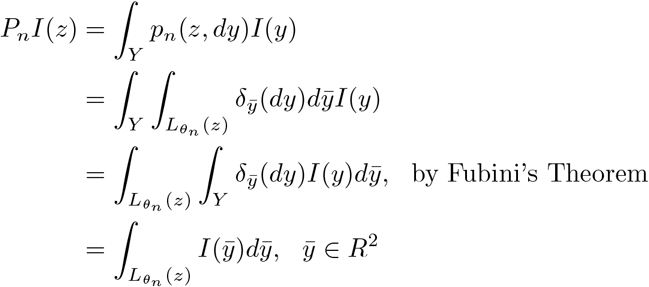

## Appendix B

### Registration Accuracy

Similar to other groups [35], we evaluated accuracy of alignment between 2D histology and 3D MRI by looking at sets of discrete points (pixels) labeled in 2D versus corresponding voxels labeled in 3D and subsequently deformed to 2D (see Figure 3). Here, our sets of points were the sets of pixels (voxels) within a particular MTL subregion as delineated on 2D histology images and 3D MRI (see Section 2.5). We restricted our attention to particular subregions of interest (amygdala, ERC, CA1, subiculum), and measured accuracy by Dice Score and 95th Percentile Hausdorff distance for each region on each slice of one brain. Average overlap scores were 0.85, 0.82, 0.74, 0.65 for amygdala, ERC, CA1, and subiculum, respectively, while average 95th percentile Hausdorff distance was 1.886 mm, 1.039 mm, 1.972 mm, and 1.746 mm for amygdala, ERC, CA1, and subiculum, respectively.

## Appendix C

### Scattering Transform

The Scattering Transform, by Mallat and Bruna [37, 38], defines a cascade of alternating non-linear and non-commuting operators that map functions *J*_*n*_(·) ∈ *L*^2^(*R*^*d*^) to representations, 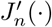. A scattering propogator takes signals down a path of alternating convolutions with wavelets (localized waveforms) and modulus operations. The path, *p*, is defined by a set of parameters *λ*∈ 2^ℤ^ that scale a mother wavelet, *ψ*, so as to capture lower and lower frequencies.

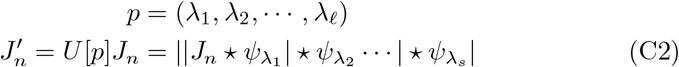

We compute a subsampled Scattering Transform 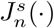 of each of our histology images, using an algorithm similar to the “Filterbank” algorithm [74, 75] in which images are downsampled in parallel with scattering.

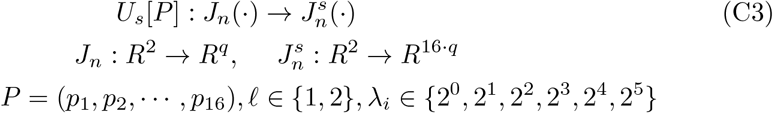

We use 16 paths, *p*_*i*_ ∈ *P* of length 1 or 2 and a high pass Gaussian filter, with width dilated according to *λ*_*i*_, in place of a traditional wavelet to achieve a representation both translation and rotation invariant in addition to Lipschitz continuous to small deformations. Each of the R,G,B channels of histology images are propagated independently along the same paths. Histology images are downsampled by a factor of 32 to reach the approximate resolution of MRI. Together, the subsampling and scattering of each channel yield a total of 48 scattering coefficients for each pixel in the downsampled histology image.

## Appendix D

### EM Algorithm for Optimization

The iterative algorithm in step C (Algorithm 2) is based on the EM algorithm implying it is monotonic in the cost. The complete-data likelihood for each histology plane *n* = 1, 2, …*N* as a function of parameters, *θ* = *ϕ*_*n*_ is

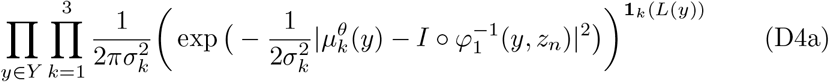

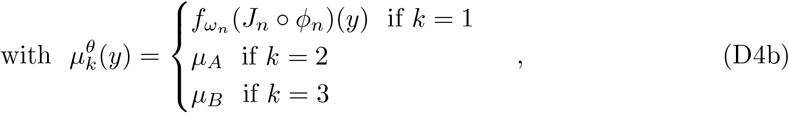

where *μ*_*A*_ and *μ*_*B*_ represent artifact and background. The E-step takes the conditional expectation of the complete data log-likelihood with respect to the incomplete data, and the previous parameters *θ*^*old*^. The *M* -step generates our sequence of parameters:

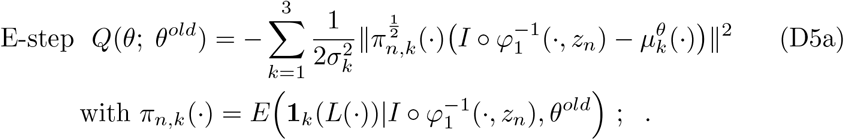

The spatial field of weights *π*_*n,k*_(·) is the conditional expectation of the indicator **1**_*k*_(*L*(·)). We implement the Generalized EM (GEM) algorithm (see [39]) solving the maximization step:

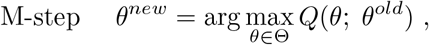

which generate a sequence with increasing log-likelihood:

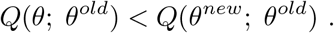

## Appendix E

### Dataset Demographics

**Table E1.** Donor demographics and pathological staging.

| Category | Subject 1 | Subject 2 |
| --- | --- | --- |
| Age | 93 | 87 |
| Gender | M | F |
| Clinical Diagnosis | Dementia | Dementia |
| Braak NFT stage | VI/VI | VI/VI |
| CERAD Neuritic<br>Plaque Score [76] | B | B |
| TDP-43 | - | + |
| Pathologic Diagnosis<br>[7] | High level AD patho-<br>logic change; multiple<br>cerebral infarcts | High level AD patho-<br>logic change |

## Appendix F

### UNET Details

To identify individual tau tangles in histology images, we trained a UNET to predict per pixel probabilities of tau and subsequently segmented probability maps into discrete “tangles” using opencv’s implementation of the watershed algorithm [64]. The architecture is summarized in table E. 10-fold cross validation was used to estimate the accuracy of UNET probability predictions prior to segmentation with the watershed algorithm. Table F3 demonstrates per-fold and average performance for the trained UNET for one of our brain samples.

**Table F2.**
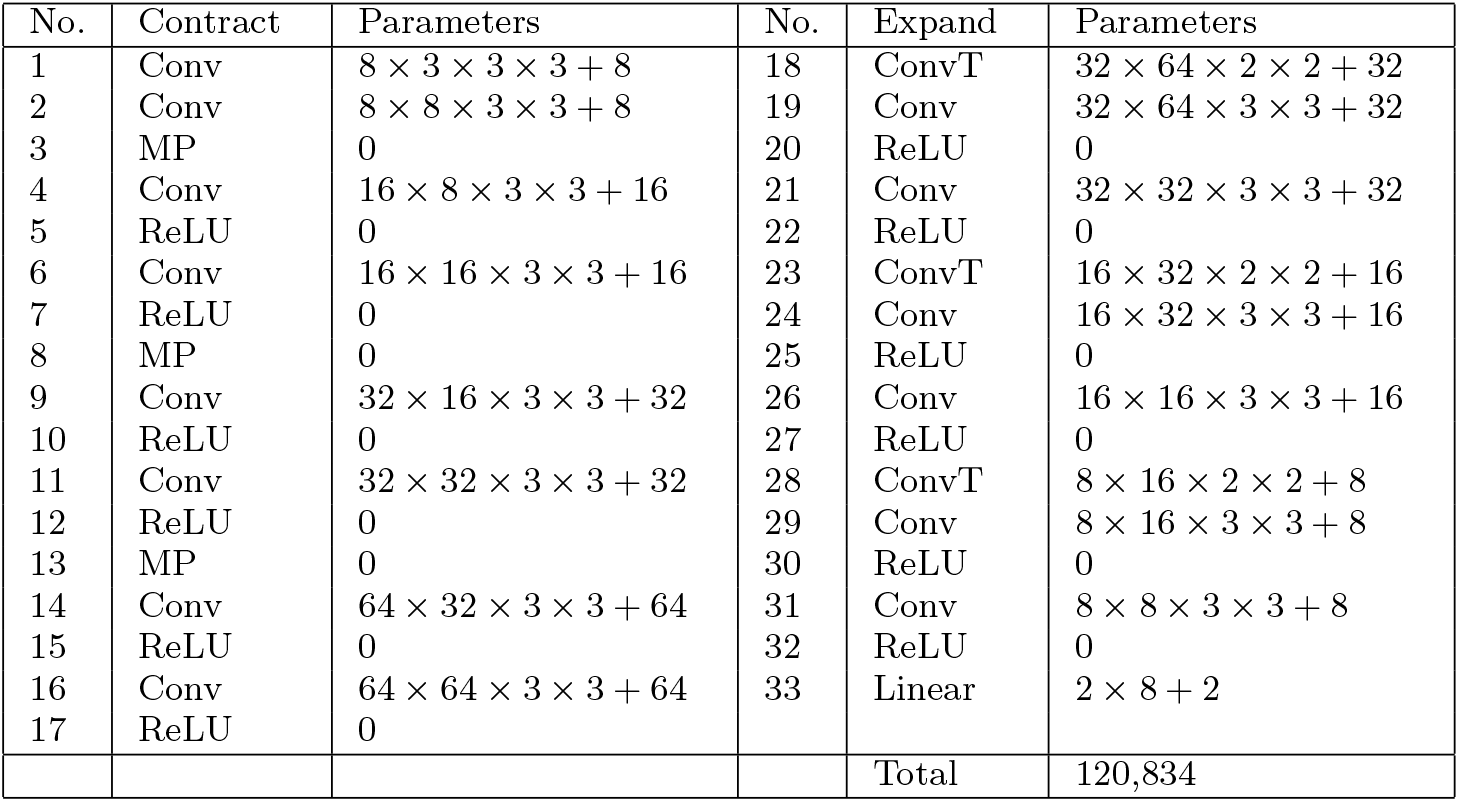
Structure of UNET trained to detect tau tangles. Contraction layers are shown in the left 3 columns, and expansion layers in the right 3 columns. Number of parameters listed correspond to linear filters + bias vector .Conv: 3 × 3 Convolution with stride 1, MP: 2 × 2 max pool, ReLU: Rectified Linear Unit, ConvT: 2× 2 transposed convolution with stride 2. Note number of features doubles in the expansion layers due to concatenation with the contraction layers (skip connections).

**Table F3.** 10-fold cross validation accuracy statistics for training data of single brain sample.

| Trial | AUC | Precision | Recall | Accuracy |
| --- | --- | --- | --- | --- |
| 1 | 0.9997 | 0.0455 | 1.0000 | 0.9928 |
| 2 | 0.9983 | 0.0938 | 0.9917 | 0.9829 |
| 3 | 0.9963 | 0.1148 | 0.9846 | 0.9706 |
| 4 | 0.9984 | 0.2079 | 0.9971 | 0.9877 |
| 5 | 0.9796 | 0.3010 | 0.9242 | 0.9543 |
| 6 | 0.9915 | 0.0500 | 0.9620 | 0.9597 |
| 7 | 0.9865 | 0.2462 | 0.9031 | 0.9899 |
| 8 | 0.9239 | 0.0081 | 0.7079 | 0.9520 |
| 9 | 0.9989 | 0.1714 | 1.000 | 0.9939 |
| 10 | 0.9867 | 0.0255 | 0.9406 | 0.9454 |
| Average | 0.9860 | 0.1264 | 0.9411 | 0.9729 |

## Appendix G

### Resampling in Mai Atlas Space

To compute and compare NFT density measures across brain samples, we rigidly aligned all samples to the reference brain in the Mai Paxinos Atlas [77]. We used a manual alignment tool, created in-house, to select optimal alignments between surface renderings of the Hippocampus, Amygdala, and ERC of our brain samples and that of the Mai brain.

Distributions of NFT density were computed in the coordinates of the Mai atlas, according to choice of *π*(*x, x*′) governing physical spatial spread, and *k*((*y, α*_*y*_),·) governing smoothing over conditional feature distributions (Equation 12b). In all cases, total mass (2D cross-sectional tissue area of histological images) and total number of NFTs were conserved. To achieve this, initial NFT feature values (counts per MTL subregion) were reformulated following physical transformation (Equation 10b) as counts of NFTs per MTL subregion per weight of particle (i.e. total cross-sectional area):

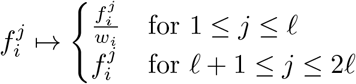

We highlight three different modes of resampling. Volumetric resampling (e.g. at mm resolution) was computed with a 3D isotropic Gaussian kernel with width, *σ*, and with new particles in a regular lattice, *x*′ ∈ *X*′.

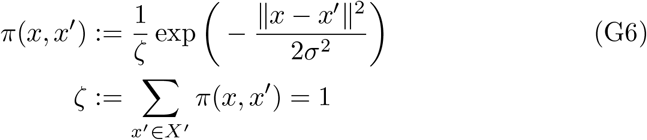

Resampling over 2D manifolds (e.g. the surface of ERC, Amygdala, CA1, or Subiculum) was computed using a nearest neighbor kernel, assigning all weight (tissue area) and NFTs from a particle at the fine scale to a single particle on the 2D manifold (e.g. vertex of a triangular mesh).

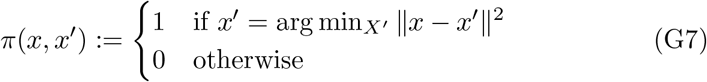

Finally, resampling to a regular 1D lattice (e.g. the rostral-caudal axis of the human brain) was computed using an anisotropic Gaussian kernel to spread particle mass widely in two dimensions and narrowly in the third, with dimensions treated independently.

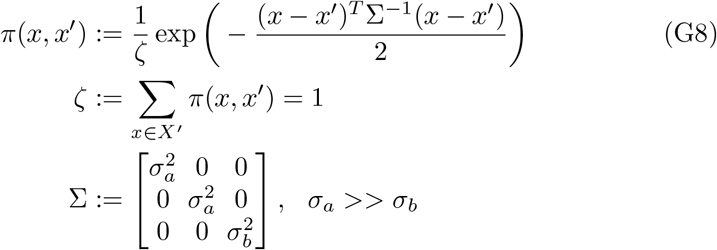

In each case, feature reduction occurred via computation of expected first moments, as described in section 2.7.

## Appendix H

### Laplace Beltrami Smoothing Solution

The variational solution to Equation 14 is given by:

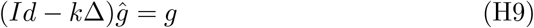

where Δ is a Laplacian operator. Here, we take Δ as the Laplace Beltrami operator and compute an eigenbasis (ℬ = {*β*_1_, …, *β*_*N*_}) and eigenvalues ({*λ*_1_, …, *λ*_*N*_}) via the Finite Elements Method (FEM) [66]. Expansion of (H9) in this eigenbasis yields smoothed *ĝ*_*a*_(·),*ĝ*_*τ*_ (·) for choice of parameter k:

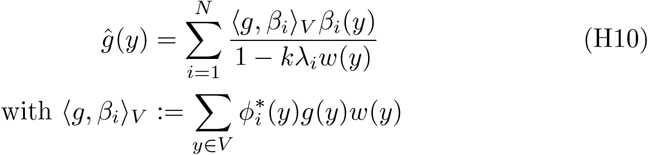

Both *ĝ*_*a*_(·) and *ĝ*_*τ*_ (·) are normalized independently so total cross sectional area and numbers of NFTs projected to the surface are conserved before and after smoothing. NFT densities are computed as the ratio of the normalized, smoothed functions: 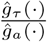 and plotted over the surfaces of given MTL subregions (see Figure 5).

## References

[1] Association, A.: 2021 Alzheimer’s disease facts and figures: Race, Ethnicity and Alzheimer’s in America. Race, Ethnicity and Alzheimer’s in America 13(3), 1–104 (2021)

[2] Sperling, R.A., Aisen, P.S., Beckett, L.A., Bennett, D.A., Craft, S., Fagan, A.M., Iwatsubo, T., Jack, C.R., Kaye, J., Montine, T.J., Park, D.C., Reiman, E.M., Rowe, C.C., Siemers, E., Stern, Y., Yaffe, K., Carrillo, M.C., Thies, B., Morrison-Bogorad, M., Wagster, M.V., Phelps, C.H.: Toward defining the preclinical stages of Alzheimer’s disease: Recommendations from the National Institute on Aging-Alzheimer’s Association workgroups on diagnostic guidelines for Alzheimer’s disease. Alzheimer’s and Dementia 7(3), 280–292 (2011). https://doi.org/10.1016/j.jalz.2011.03.003

[3] Nauen, D.W., Troncoso, J.C.: Amyloid-beta is present in human lymph nodes and greatly enriched in those of the cervical region. Alzheimer’s and Dementia (April), 1–6 (2021). https://doi.org/10.1002/alz.12385

[4] Villemagne, V.L., Doré, V., Burnham, S.C., Masters, C.L., Rowe, C.C.: Imaging tau and amyloid-? proteinopathies in Alzheimer disease and other conditions. Nature Reviews Neurology 14(4), 225–236 (2018). https://doi.org/10.1038/nrneurol.2018.9

[5] Braak, H., Braak, E.: Neuropathological stageing of Alzheimer-related changes. Acta Neuropathol. 82(4), 239–259 (1991). https://doi.org/10.1007/BF00308809

[6] Braak, H., Alafuzoff, I., Arzberger, T., Kretzschmar, H., Tredici, K.: Staging of Alzheimer disease-associated neurofibrillary pathology using paraffin sections and immunocytochemistry. Acta Neuropathologica 112(4), 389–404 (2006). https://doi.org/10.1007/s00401-006-0127-z

[7] Hyman, B.T., Phelps, C.H., Beach, T.G., Bigio, E.H., Cairns, N.J., Carrillo, M.C., Dickson, D.W., Duyckaerts, C., Frosch, M.P., Masliah, E., Mirra, S.S., Nelson, P.T., Schneider, J.A., Thal, D.R., Thies, B., Trojanowski, J.Q., Vinters, H.V., Montine, T.J.: National Institute on Aging-Alzheimer’s Association guidelines for the neuropathologic assessment of Alzheimer’s disease. Alzheimer’s and Dementia 8(1), 1–13 (2012). https://doi.org/10.1016/J.JALZ.2011.10.007

[8] Jack, C.R., Bennett, D.A., Blennow, K., Carrillo, M.C., Dunn, B., Haeberlein, S.B., Holtzman, D.M., Jagust, W., Jessen, F., Karlawish, J., Liu, E., Molinuevo, J.L., Montine, T., Phelps, C., Rankin, K.P., Rowe, C.C., Scheltens, P., Siemers, E., Snyder, H.M., Sperling, R., Elliott, C., Masliah, E., Ryan, L., Silverberg, N.: NIA-AA Research Framework: Toward a biological definition of Alzheimer’s disease. Alzheimer’s Dement. 14(4), 535–562 (2018). https://doi.org/10.1016/j.jalz.2018.02.018

[9] Zetterberg, H., Bendlin, B.B.: Biomarkers for Alzheimer’s disease—preparing for a new era of disease-modifying therapies. Molecular Psychiatry 26(1), 296–308 (2021). https://doi.org/10.1038/s41380-020-0721-9

[10] Young, P.N.E., Estarellas, M., Coomans, E., Srikrishna, M., Beaumont, H., Maass, A., Venkataraman, A.V., Lissaman, R., Jiménez, D., Betts, M.J., McGlinchey, E., Berron, D., O’Connor, A., Fox, N.C., Pereira, J.B., Jagust, W., Carter, S.F., Paterson, R.W., Schöll, M.: Imaging biomarkers in neurodegeneration: Current and future practices. Alzheimer’s Research and Therapy 12(1) (2020). https://doi.org/10.1186/s13195-020-00612-7

[11] Younes, L., Albert, M., Miller, M.I.: Inferring changepoint times of medial temporal lobe morphometric change in preclinical Alzheimer’s disease. NeuroImage: Clinical 5, 178–187 (2014). https://doi.org/10.1016/j.nicl.2014.04.009

[12] Younes, L., Albert, M., Moghekar, A., Soldan, A., Pettigrew, C., Miller, M.I.: Identifying changepoints in biomarkers during the preclinical phase of Alzheimer’s disease. Frontiers in Aging Neuroscience 11(APR) (2019). https://doi.org/10.3389/fnagi.2019.00074

[13] Kulason, S., Tward, D.J., Brown, T., Sicat, C.S., Liu, C.F., Ratnanather, J.T., Younes, L., Bakker, A., Gallagher, M., Albert, M., Miller, M.I.: Cortical thickness atrophy in the transentorhinal cortex in mild cognitive impairment. NeuroImage: Clin. 21, 101617 (2019). https://doi.org/10.1016/j.nicl.2018.101617

[14] Lee, B.C., Tward, D.J., Mitra, P.P., Miller, M.I.: On variational solutions for whole brain serial-section histology using a Sobolev prior in the computational anatomy random orbit model. PLoS Computational Biology 14(12), 1–20 (2018). https://doi.org/10.1371/journal.pcbi.1006610

[15] Tward, D., Li, X., Huo, B., Lee, B., Mitra, P., Miller, M.: 3D Mapping of Serial Histology Sections with Anomalies Using a Novel Robust Deformable Registration Algorithm 3D Mapping of Serial Sections via Robust Deformable Registration 163. In: LNCS, vol. 11846, pp. 162–173 (2019). https://doi.org/10.1007/978-3-030-33226-618

[16] Lee, B.C., Lin, M.K., Fu, Y., Hata, J., Miller, M.I., Mitra, P.P.: Multimodal cross-registration and quantification of metric distortions in marmoset whole brain histology using diffeomorphic mappings. Journal of Comparative Neurology 529(2), 281–295 (2021) https://arxiv.org/abs/ 1805.04975v2. https://doi.org/10.1002/cne.24946

[17] Iglesias, J.E., Insausti, R., Lerma-Usabiaga, G., Bocchetta, M., Van Leemput, K., Greve, D.N., van der Kouwe, A., Fischl, B., Caballero-Gaudes, C., Paz-Alonso, P.M.: A probabilistic atlas of the human thalamic nuclei combining ex vivo MRI and histology. NeuroImage 183(July), 314– 326 (2018) https://arxiv.org/abs/1806.08634. https://doi.org/10.1016/j.neuroimage.2018.08.012

[18] Perens, J., Salinas, C.G., Skytte, J.L., Roostalu, U., Dahl, A.B., Dyrby, T.B., Wichern, F., Barkholt, P., Vrang, N., Jelsing, J., Hecksher-Sørensen, J.: An Optimized Mouse Brain Atlas for Automated Mapping and Quantification of Neuronal Activity Using iDISCO+ and Light Sheet Fluorescence Microscopy. Neuroinformatics 19(3), 433–446 (2021). https://doi.org/10.1007/s12021-020-09490-8

[19] Hillman, E.M.C., Voleti, V., Li, W., Yu, H.: Light-Sheet Microscopy in Neuroscience. Annual Review of Neuroscience 42, 295–313 (2019). https://doi.org/10.1146/annurev-neuro-070918-050357

[20] Bergenstråhle, J., Larsson, L., Lundeberg, J.: Seamless integration of image and molecular analysis for spatial transcriptomics workflows. BMC Genomics 21(1), 1–7 (2020). https://doi.org/10.1186/s12864-020-06832-3

[21] Chen, W.T., Lu, A., Craessaerts, K., Pavie, B., Sala Frigerio, C., Corthout, N., Qian, X., Laláková, J., Kühnemund, M., Voytyuk, I., Wolfs, L., Mancuso, R., Salta, E., Balusu, S., Snellinx, A., Munck, S., Jurek, A., Fernandez Navarro, J., Saido, T.C., Huitinga, I., Lundeberg, J., Fiers, M., De Strooper, B.: Spatial Transcriptomics and In Situ Sequencing to Study Alzheimer’s Disease. Cell 182(4), 976–99119 (2020). https://doi.org/10.1016/j.cell.2020.06.038

[22] Miller, M.I., Fan, J., Tward, D.J.: Multi scale diffeomorphic metric mapping of spatial transcriptomics datasets. In: IEEE Computer Society Conference on Computer Vision and Pattern Recognition Workshops, pp. 4467–4475. IEEE, ??? (2021). https://doi.org/10.1109/CVPRW53098.2021.00504

[23] Palla, G., Fischer, D.S., Regev, A., Theis, F.J.: Spatial components of molecular tissue biology. Nature Biotechnology (2022). https://doi.org/10.1038/s41587-021-01182-1

[24] Grenander, U., Miller, M.I.: Computational Anatomy: An Emerging Discipline. Applied Mathematics 56(4), 617–694 (1998)

[25] Grenander, U., Miller, M.I.: Pattern Theory: From Representation to Inference. OUP Oxford, ??? (2006). https://books.google.com/books?id=rQlREAAAQBAJ

[26] Miller, M.I., Younes, L., Trouvé, A.: Diffeomorphometry and geodesic positioning systems for human anatomy. TECHNOLOGY 02(01), 36–43 (2014). https://doi.org/10.1142/s2339547814500010

[27] Miller, M.I., Arguillére, S., Tward, D.J., Younes, L.: Computational anatomy and diffeomorphometry: A dynamical systems model of neuroanatomy in the soft condensed matter continuum. Wiley Interdisciplinary Reviews: Systems Biology and Medicine 10(6), 1–42 (2018). https://doi.org/10.1002/wsbm.1425

[28] Miller, M.I., Younes, L.: Group actions, homeomorphisms, and matching: A general framework. International Journal of Computer Vision 41, 61–84 (2001)

[29] Miller, M.I., Trouvé, A., Younes, L.: On the metrics and Euler-Lagrange equations of computational anatomy. Annual Review of Biomedical Engineering 4, 375–405 (2002). https://doi.org/10.1146/annurev.bioeng.4.092101.125733

[30] Avants, B.B., Epstein, C.L., Grossman, M., Gee, J.C.: Symmetric diffeomorphic image registration with cross-correlation: Evaluating automated labeling of elderly and neurodegenerative brain. Medical Image Analysis 12(1), 26–41 (2008). https://doi.org/10.1016/j.media.2007.06.004

[31] Christensen, G.E., Johnson, H.J.: Consistent image registration. IEEE Transactions on Medical Imaging 20(7), 568–582 (2001). https://doi.org/10.1109/42.932742

[32] Avants, B.B., Epstein, C.L., Grossman, M., Gee, J.C.: Symmetric Diffeomorphic Image Registration with Cross-Correlation: Evaluating Automated Labeling of Elderly and Neurodegenerative Brain

[33] Pluim, J.P.W., Maintz, J.B.A.A., Viergever, M.A.: Mutual-information-based registration of medical images: A survey. IEEE Transactions on Medical Imaging 22(8), 986–1004 (2003). https://doi.org/10.1109/TMI.2003.815867

[34] Heinrich, M.P., Jenkinson, M., Bhushan, M., Matin, T., Gleeson, F.V., Brady, S.M., Schnabel, J.A.: MIND: Modality independent neighbourhood descriptor for multi-modal deformable registration. Medical Image Analysis 16(7), 1423–1435 (2012). https://doi.org/10.1016/j.media.2012.05.008

[35] Yushkevich, P.A., de Onzoño Martin, M.M.I., Ittyerah, R., Lim, S., Lavery, M., Wang, J., Hung, L.Y., Vergnet, N., Ravikumar, S., Xie, L., Dong, M., DeFlores, R., Cui, S., McCollum, L., Ohm, D.T., Robinson, J.L., Schuck, T., Grossman, M., Tisdall, M.D., Prabhakaran, K., Mizsei, G., Das, S.R., Artacho-Pérula, E., del Mar Arroyo Jiménez, M., López, M.M., Rabal, M.P.M., Romero, F.J.M., Lee, E.B., Trojanowski, J.Q., Wisse, L.E.M., Wolk, D.A., Irwin, D.J., Insausti, R.: 3d mapping of tau neurofibrillary tangle pathology in the human medial temporal lobe. In: 2020 IEEE 17th International Symposium on Biomedical Imaging (ISBI), pp. 1312–1316 (2020). https://doi.org/10.1109/ISBI45749.2020.9098462

[36] Tward, D., Brown, T., Kageyama, Y., Patel, J., Hou, Z., Mori, S., Albert, M., Troncoso, J., Miller, M.: Diffeomorphic Registration With Intensity Transformation and Missing Data: Application to 3D Digital Pathology of Alzheimer’s Disease. Front. Neurosci. 14(February), 1–18 (2020). https://doi.org/10.3389/fnins.2020.00052

[37] Mallat, S.: Group invariant scattering. Commun. Pur. Appl. Math. 65(10), 1331–1398 (2012)

[38] Bruna, J., Mallat, S.: Invariant scattering convolution networks. IEEE Trans. Pattern Anal. Mach. Intell. 35(8), 1872–1886 (2013) https://arxiv.org/abs/1203.1513. https://doi.org/10.1109/TPAMI.2012.230

[39] Dempster, A.P., Laird, N.M., Rubin, D.B.: Maximum likelihood from incomplete data via the em algorithm. J. R. Stat. Soc. Series B 39(1), 1–38 (1977)

[40] Miller, M.I., Tward, D., Trouvé, A.: Hierarchical Computational Anatomy: Unifying the Molecular to Tissue Continuum via Measure Representations of the Brain. bioRxiv (2021). https://doi.org/10.1101/2021.04.19.440540

[41] Tournier, J.-D., Mori, S., Leemans, A., Morgan, R.H., Reson, M., Author, M.: Diffusion Tensor Imaging and Beyond NIH Public Access Author Manuscript. Magn Reson Med 65(6), 1532–1556 (2011). https://doi.org/10.1002/mrm.22924.Diffusion

[42] Bracewell, R.N.: Strip Integration in Radio Astronomy. Australian Journal of Physics 9, 198 (1956). https://doi.org/10.1071/PH560198

[43] Zhao, S.-R., Halling, H.: A new fourier method for fan beam reconstruction. In: 1995 IEEE Nuclear Science Symposium and Medical Imaging Conference Record, vol. 2, pp. 1287–12912 (1995). https://doi.org/10.1109/NSSMIC.1995.510494

[44] Dupuis, P., Grenander, U., Miller, M.I.: Variational problems on flows of diffeomorphisms for image matching. Quarterly of Applied Mathematics 56(3), 587–600 (1998)

[45] Joshi, S., Miller, M.I.: Maximum a posteriori estimation with Good’s roughness for three-dimensional optical-sectioning microscopy. J. Opt. Soc. Am. A 10(5), 1078–1085 (1993). https://doi.org/10.1364/JOSAA.10.001078

[46] Gibson, S.F., Lanni, F.: Diffraction by a circular aperture as a model for three-dimensional optical microscopy. J. Opt. Soc. Am. A 6(9), 1357–1367 (1989). https://doi.org/10.1364/JOSAA.6.001357

[47] Snyder, D., Thomas Jr., L., Ter-Pogossian, M.: Mathematical model for positron-emission tomography systems having time-of-flight measurements. IEEE Transactions on Nuclear Science NS-28(3), 3575–83 (1981)

[48] Barrett, H.H.: Iii the radon transform and its applications. Progress in Optics, vol. 21, pp. 217–286. Elsevier (1984). https://doi.org/10.1016/S0079-6638(08)70123-9. https://www.sciencedirect.com/science/article/pii/S0079663808701239

[49] Snyder, D., Cox, J.: An overview of reconstruction tomography and limitations imposed by a finite number of projections. In: Proceedings of Workshop on Reconstruction Tomography in Diagnostic Radiology and Nuclear Medicine, Puerto Rico (1975)

[50] Beg, M.F., Miller, M.I., Trouvé, A., Younes, L.: Computing large deformation metric mappings via geodesic flows of diffeomorphisms. Int. J. Comput. Vis. 61(2), 139–157 (2005). https://doi.org/10.1023/B:VISI.0000043755.93987.aa

[51] Tward, D.J.: An optical flow based left-invariant metric for natural gradient descent in affine image registration. Frontiers in Applied Mathematics and Statistics, 61

[52] CIBC Seg3D: Volumetric Image Segmentation and Visualization. Scientific Computing and Imaging Institute (SCI), Download from: http://www.seg3d.org (2016)

[53] Berron, D., Vieweg, P., Hochkeppler, A., Pluta, J.B., Ding, S.L., Maass, A., Luther, A., Xie, L., Das, S.R., Wolk, D.A., Wolbers, T., Yushkevich, P.A., Düzel, E., Wisse, L.E.M.: A protocol for manual segmentation of medial temporal lobe subregions in 7 Tesla MRI. NeuroImage: Clinical 15(May), 466–482 (2017). https://doi.org/10.1016/j.nicl.2017.05.022

[54] Wisse, L.E.M., Gerritsen, L., Zwanenburg, J.J.M., Kuijf, H.J., Luijten, P.R., Biessels, G.J., Geerlings, M.I.: Subfields of the hippocampal formation at 7t mri: In vivo volumetric assessment. NeuroImage 61(4), 1043–1049 (2012). https://doi.org/10.1016/j.neuroimage.2012.03.023

[55] Yushkevich, P., Avants, B., Pluta, J., Das, S., Minkoff, D., Mechanichamilton, D., Glynn, S., Pickup, S., Liu, W., Gee, J.: A high-resolution computational atlas of the human hippocampus from postmortem magnetic resonance imaging at 9.4 t. NeuroImage 44(2), 385– 398 (2009). https://doi.org/10.1016/j.neuroimage.2008.08.042

[56] Ding, S.-L.: Comparative anatomy of the prosubiculum, subiculum, presubiculum, postsubiculum, and parasubiculum in human, monkey, and rodent. Journal of Comparative Neurology 521(18), 4145– 4162 (2013) https://arxiv.org/abs/https://onlinelibrary.wiley.com/doi/pdf/10.1002/cne.23416.https://doi.org/10.1002/cne.23416

[57] Ding, S.-L., Hoesen, G.: Organization and detailed parcellation of human hippocampal head and body regions based on a combined analysis of cyto-and chemoarchitecture. J Comp Neurol 523, 2233–2253 (2015). https://doi.org/10.1002/cne.23786

[58] Insausti, R., Tuñón, T., Sobreviela, T., Insausti, A.M., Gonzalo, L.M.: The human entorhinal cortex: A cytoarchitectonic analysis. Journal of Comparative Neurology 355(2), 171–198 (1995) https://arxiv.org/abs/https://onlinelibrary.wiley.com/doi/pdf/10.1002/cne.903550203.https://doi.org/10.1002/cne.903550203

[59] Insausti, R., Córcoles-Parada, M., Ubero, M.M., Rodado, A., Insausti, A.M., Muñoz-López, M.: Cytoarchitectonic areas of the Gyrus ambiens in the human brain. Frontiers in Neuroanatomy 13 (2019). https://doi.org/10.3389/fnana.2019.00021

[60] Olga, K., Zilles, K., Palomero-Gallagher, N., Schleicher, A., Mohlberg, H., Bludau, S., Amunts, K.: Receptor-driven, multimodal mapping of the human amygdala. Brain Structure and Function 223 (2018). https://doi.org/10.1007/s00429-017-1577-x

[61] Amunts, K., Olga, K., Kindler, M., Pieperhoff, P., Mohlberg, H., Shah, N., Habel, U., Schneider, F., Zilles, K.: Cytoarchitectonic mapping of the human amygdala, hippocampal region and entorhinal cortex: intersubject variability and probability maps. Anatomy and embryology 210, 343–52 (2006). https://doi.org/10.1007/s00429-005-0025-5

[62] Stouffer, K.M., Wang, Z., Xu, E., Lee, K., Lee, P., Miller, M.I., Tward, D.J.: From picoscale pathology to decascale disease: Image registration with a scattering transform and varifolds for manipulating multiscale data. In: Syeda-Mahmood, T., Li, X., Madabhushi, A., Greenspan, H., Li, Q., Leahy, R., Dong, B., Wang, H. (eds.) Multimodal Learning for Clinical Decision Support, pp. 1–11. Springer, Cham (2021)

[63] Ronneberger, O., Fischer, P., Brox, T.: U-Net: Convolutional Networks for Biomedical Image Segmentation. In: International Conference on Medical Image Computing and Computer-assisted Intervention, pp. 234–241. Springer, ??? (2015)

[64] Bradski, G.: The OpenCV Library. Dr. Dobb’s J. Softw. Tools (2000)

[65] Otsu, N.: A threshold selection method from gray-level histograms. IEEE Transactions on Systems, Man, and Cybernetics 9(1), 62–66 (1979). https://doi.org/10.1109/TSMC.1979.4310076

[66] Qiu, A., Bitouk, D., Miller, M.I.: Smooth functional and structural maps on the neocortex via orthonormal bases of the laplace-beltrami operator. IEEE Transactions on Medical Imaging 25(10), 1296–1306 (2006). https://doi.org/10.1109/TMI.2006.882143

[67] Poulin, S.P., Dautoff, R., Morris, J.C., Barrett, L.F., Dickerson, B.C.: Amygdala atrophy is prominent in early Alzheimer’s disease and relates to symptom severity. Psychiatry Research - Neuroimaging 194(1), 7–13 (2011). https://doi.org/10.1016/j.pscychresns.2011.06.014

[68] Ortner, M., Pasquini, L., Barat, M., Alexopoulos, P., Grimmer, T., Förster, S., Diehl-Schmid, J., Kurz, A., Förstl, H., Zimmer, C., Wohlschläger, A., Sorg, C., Peters, H.: Progressively disrupted intrinsic functional connectivity of basolateral amygdala in very early Alzheimer’s disease. Frontiers in Neurology 7(SEP), 1–9 (2016). https://doi.org/10.3389/fneur.2016.00132

[69] Nelson, P.T., Abner, E.L., Patel, E., Anderson, S., Wilcock, D.M., Kryscio, R.J., Van Eldik, L.J., Jicha, G.A., Gal, Z., Nelson, R.S., Nelson, B.G., Gal, J., Azam, M.T., Fardo, D.W., Cykowski, M.D.: The amygdala as a locus of pathologic misfolding in neurodegenerative diseases. Journal of Neuropathology and Experimental Neurology 77(1), 2–20 (2018). https://doi.org/10.1093/jnen/nlx099

[70] Miller, M.I., Ratnanather, J.T., Tward, D.J., Brown, T., Lee, D.S., Ketcha, M., Mori, K., Wang, M.C., Mori, S., Albert, M.S., Younes, L., Rodzon, B., Selnes, O., Gottesman, R., Sacktor, N., McKhann, G., Turner, S., Farrington, L., Grega, M., D’Agostino, D., Feagen, S., Dolan, D., Dolan, H., Kolasny, A., Schneider, W., O’Brien, R., Moghekar, A., Meehan, R., Scherer, R., Meinert, C., Shade, D., Ervin, A., Jones, J., Toepfner, M., Parlett, L., Patterson, A., Lassiter, L., Li, S., Lu, Y., Troncoso, J., Crain, B., Pletnikova, O., Rudow, G., Fisher, K.: Network neurodegeneration in Alzheimer’s disease via MRI based shape diffeomorphometry and high-field atlasing. Frontiers in Bioengineering and Biotechnology 3(MAY), 1–16 (2015). https://doi.org/10.3389/fbioe.2015.00054

[71] Miller, M.I., Younes, L., Ratnanather, J.T., Brown, T., Trinh, H., Lee, D.S., Tward, D., Mahon, P., Mori, S., Albert, M., Team, R.: Amygdalar Atrophy in Symptomatic AD Based on Diffeomorphometry: The BIOCARD Cohort. Neurobiol Aging 36(1), 3–10 (2015). https://doi.org/10.1016/j.neurobiolaging.2014.06.032

[72] Johansson, M., Stomrud, E., Insel, P.S., Leuzy, A., Johansson, P.M., Smith, R., Ismail, Z., Janelidze, S., Palmqvist, S., van Westen, D., Mattsson-Carlgren, N., Hansson, O.: Mild behavioral impairment and its relation to tau pathology in preclinical Alzheimer’s disease. Translational Psychiatry 11(1) (2021). https://doi.org/10.1038/s41398-021-01206-z

[73] Matuskova, V., Ismail, Z., Nikolai, T., Markova, H., Cechova, K., Nedelska, Z., Laczo, J., Wang, M., Hort, J., Vyhnalek, M.: Mild Behavioral Impairment Is Associated With Atrophy of Entorhinal Cortex and Hippocampus in a Memory Clinic Cohort. Frontiers in Aging Neuroscience 13(May), 1–9 (2021). https://doi.org/10.3389/fnagi.2021.643271

[74] Mallat, S.: Recursive interferometric representations. In: European Signal Processing Conference, pp. 716–720 (2010)

[75] SIfre, L., Mallat, S.: Rigid-Motion Scattering for Texture Classification (2014)

[76] Mirra, S.S., Heyman, A., McKeel, D., Sumi, S.M., Crain, B.J., Brownlee, L.M., Vogel, F.S., Hughes, J.P., Belle, G.v., Berg, L., participating CERAD neuropathologists: The consortium to establish a registry for alzheimer’s disease (cerad). Neurology 41(4), 479–479 (1991) https://arxiv.org/abs/https://n.neurology.org/content/41/4/479.full.pdf.https://doi.org/10.1212/WNL.41.4.479

[77] Mai, J.K., Paxinos, G., Voss, T.: Atlas of the Human Brain, 3rd edn. Elsevier Inc, New York (2008)

